# Circadian control of the secretory pathway is a central mechanism in tissue homeostasis

**DOI:** 10.1101/304014

**Authors:** Joan Chang, Richa Garva, Adam Pickard, Ching-Yan Chloé Yeung, Venkatesh Mallikarjun, Joe Swift, David F. Holmes, Ben Calverley, Yinhui Lu, Antony Adamson, Helena Raymond-Hayling, Oliver Jensen, Tom Shearer, Qing Jun Meng, Karl E. Kadler

## Abstract

Collagen is the most abundant secreted protein in vertebrates that persists throughout life without renewal. The unchanging nature of collagen contrasts with observed continued collagen synthesis throughout adulthood and with conventional transcriptional and translational homeostatic mechanisms that replace damaged proteins with new copies. Here we show circadian clock regulation of procollagen transport from ER-to-Golgi and Golgi-to-plasma membrane by sequential rhythmic expression of SEC61, TANGO1, PDE4D and VPS33B. The result is nocturnal procollagen synthesis and daytime collagen fibril assembly in mice. Rhythmic collagen degradation by CTSK maintains collagen homeostasis. This circadian cycle of collagen synthesis, assembly and degradation affects only a pool of newly-synthesized collagen whilst maintaining the persistent collagen network. Disabling the circadian clock causes collagen accumulation and abnormal fibrils *in vivo*. In conclusion, our study has identified a circadian clock mechanism of protein homeostasis in which a sacrificial pool of collagen is synthesized and removed to maintain tissue function.

---

One-third of the eukaryote proteome enters the secretory pathway ^1^, including the collagens that assemble into centimeter-long fibrils in the extracellular matrix (ECM) ^2^. The fibrils account for one-third of the mass of vertebrates ^3^ and are sites of attachment for a wide range of macromolecules including integrins, making them essential for metazoan development ^4,5^. The importance of tight cellular control of collagen fibril homeostasis is exemplified in fibrosis, which is characterized by accumulation of collagen fibrils and is a feature of 45% of all deaths ^6^ including those caused by cancer ^7^, and in musculoskeletal disease in which loss of collagen leads to tissue frailty. A remarkable feature of collagen fibrils is that they are formed during embryogenesis ^8^ and remain without turnover for the life of the animal as evidenced by studies showing that the half-life of collagen in tendon and cartilage exceeds 200 years ^9–13^. This has led to the notion that collagen fibrils are static and unchanging.

However, the difficulty with the zero-turnover description of collagen is that it ignores the observation that the fibrils do not suffer from fatigue failure, which would be expected in the face of lifelong cyclic loading. In contrast to the evidence of zero replacement, fibroblasts synthesize new collagen ^14,15^ in response to mechanical and hormonal stimuli ^16–18^, and microdialysis of human Achilles tendon shows elevated levels of the C-propeptides of procollagen-I (PC-I, the precursor of collagen-I) after moderate exercise ^19,20^.

These opposing observations led to the alternative hypothesis presented in this study, in which zero turnover and continued synthesis can co-exist. We hypothesized that a pool of ‘persistent’ collagen co-exists with a pool of ‘sacrificial’ collagen, in which the latter is synthesized and removed on a daily basis under the control of the circadian clock. Support for this alternative hypothesis comes from observations of a circadian oscillation in the serum concentrations of the N- and C-propeptides of PC-I ^21,22^ and of collagen degradation products in bone ^23^. However, despite physiological and clinical observations, direct mechanistic support for these observations was lacking. Whilst the suprachiasmatic nucleus of the hypothalamus is the master circadian pacemaker, almost all tissues have self-sustaining circadian pacemakers that synchronize rhythmic tissue-specific gene expression in anticipation of environmental light and dark cycles ^24,25^. Disruption of the circadian clock leads to musculoskeletal abnormalities involving the ECM e.g. chondrocyte-specific disruption of the circadian clock resulted in progressive degeneration of the ECM in articular cartilage ^26^ and fibrosis in intervertebral disc ^27^, and mice with a global knockout (KO) of *Bmal1* ^28^ or the *ClockΔ19* mutation ^29^ develop thickened and calcified tendons with associated immobilization. These observations were indicative of circadian control of ECM homeostasis.

Here, we performed time series electron microscopy, transcriptomics and proteomics over day/night cycles, which showed that the synthesis and transport of procollagen by the protein secretory pathway in fibroblasts is regulated by the circadian clock. We showed that SEC61, TANGO1, PDE4D and VPS33B regulate collagen secretion, are 24-hour rhythmic, and are located at the entry and exit points of the endoplasmic reticulum (ER), Golgi and post-Golgi compartments, respectively. CTSK is a collagen-degrading proteinase that is rhythmic in phase with collagen degradation to maintain collagen homeostasis. The result is nocturnal PC-I synthesis and a daily wave of collagen-I with no net change in total collagen content of the tissue. Importantly, the levels of SEC61, TANGO1, PDE4D and VPS33B do not fall to zero; therefore, there is a low-level persistent secretion of collagen which has been previously detected using radioactive amino acid biosynthetic labeling of PC-I (*e.g.* ^30^). Crucially, we discovered that arrhythmic *ClockΔ19* and Scleraxis-Cre-dependent *Bmal1* deletion mutant mice accumulate collagen and have a disorganized and structurally abnormal collagen matrix that is mechanically abnormal, which demonstrates the importance of secretory pathway rhythmicity in maintaining collagen homeostasis.

## Results

### Collagen fibril network in tendon follows a rhythmic pattern during 24 hours

Tendon is a peripheral circadian clock tissue with collagen fibrils arranged parallel to the long axis of the tissue. Time-series transmission electron microscopy (TEM) of wild-type C57/BL6 mice identified a trimodal distribution of diameters (D1, <75 nm; D2, ∼75-150 nm; D3 >150 nm) that varied with time of day (Figure 1A-C and Figure S1A). The relative proportions of D1 and D2 fibrils were antiphase to that of D3 fibrils during 24 hours (Figure 1D). This was the first evidence, to our knowledge, that the distribution of collagen fibril diameters is rhythmic during 24 hours. In further experiments, we mechanically dissociated whole tendons and performed time-series mass-mapping scanning transmission electron microscopy on the dispersed fibrils (Figure S1B, C), using previously described methods ^31^. Analysis of the mass per unit length data showed an abundance of D1 fibrils from tendons sampled at early night (Figure 1E), which corresponded to the time of day when levels of D3 fibrils were building up in undisrupted tendons. Thus, D3 fibrils can be separated into D1 fibrils by mechanical disruption. Next, we examined tendon tissue by serial block face-scanning electron microscopy (SBF-SEM) (Figure 1F). Stepping through the image stacks showed bundles of collagen fibrils in near parallel alignment with the long axis of the tendon (Video 1). Automated segmentation of the fibrils in three-dimensional reconstructions showed that D2 and D3 fibrils maintained relative positions to each other (Figure 1G, Video 2). In contrast, the narrow D1 fibrils exchanged relative positions with each other and with D2 and D3 fibrils, and made contacts with different D2 and D3 fibrils (Figure 1H, Video 3). The tips of D1 fibrils were attached to the shafts of D3 fibrils, and D3 fibrils appeared to become loose and give rise to new D1 fibrils (Video 4). We also showed that fibrils sampled at ZT15 (zeitgeber time, 3 hours into darkness) had more circular outlines than those sampled at ZT3 (3 hours into light, Figure 1I). In further experiments we showed that the percentage energy loss during dynamic cyclic loading at various strain rates varied between day and night; tendons at ZT15 exhibited more energy loss than at ZT3 independent of strain rate (Figure S1D-G). Three distinct time constants were calculated from stress-relaxation curves, all of which were higher on average in ZT3 tendons than in ZT15 tendons (T_2_, in particular, was statistically significantly higher) (Figure S1F), suggesting that the tendons relax more quickly at night.

**Figure 1.**
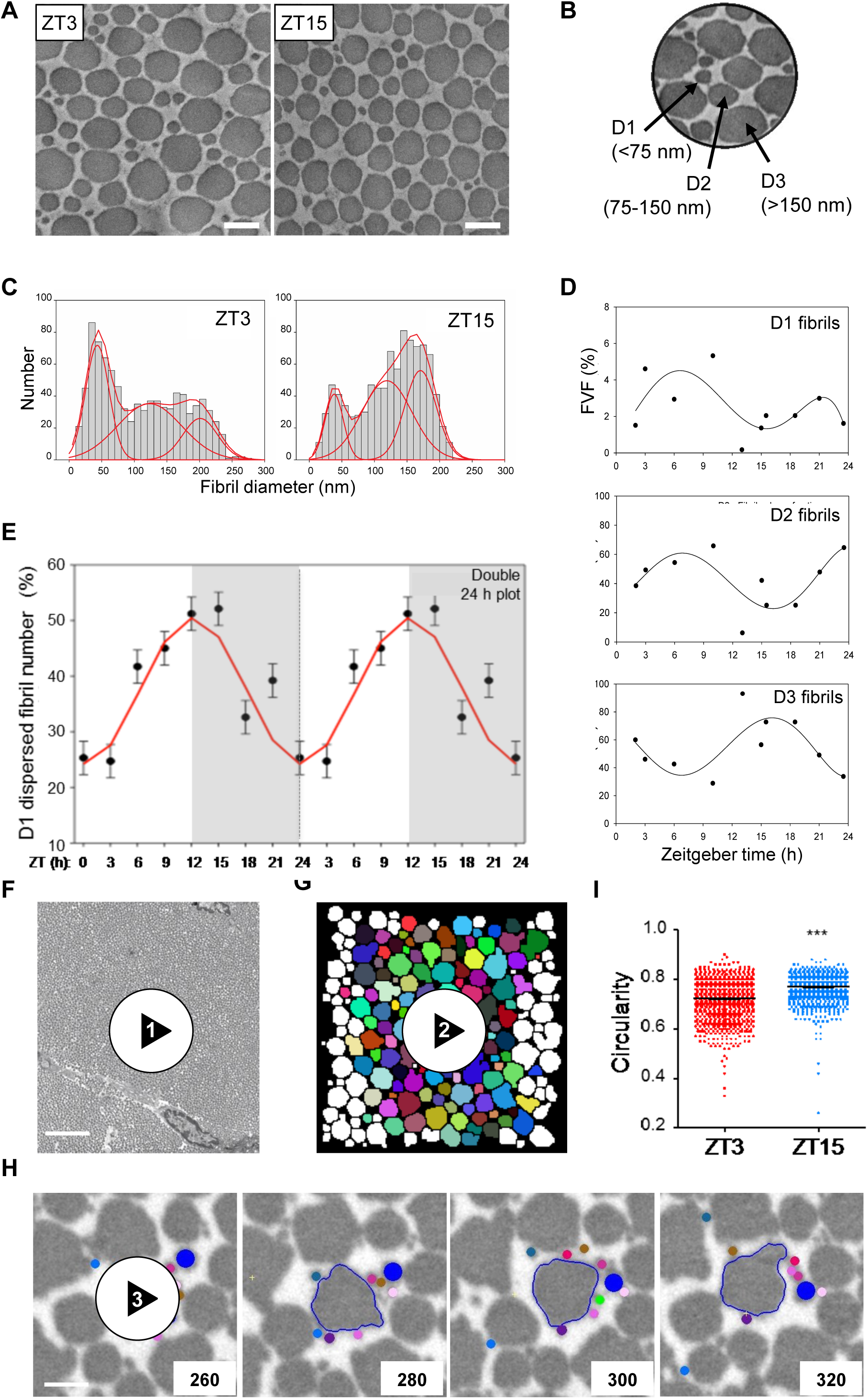
Circadian regulation of the tendon collagen extracellular matrix. **(A)** Representative transverse TEM images of Achilles tendon at ZT3 and ZT15. Bars 200 nm. **(B)** Transverse section of Achilles tendon showing characteristic D1, D2 and D3 fibrils. **(C)** Trimodal distribution of fibril diameters D1 (<75 nm), D2 (∼75-150 nm) and D3 (>150 nm). Fibril diameter distributions measured from TEM images of transverse sections of Achilles tendon sampled at ZT3 and ZT15 show a shift in collagen fibril diameter distribution (>1000 fibrils measured). **(D)** Circadian variation of D1, D2 and D3 fibrils in *in vivo* tendon derived from 3-Gaussian fitting as shown in Fig. S1. The plots show the changing fibril volume fractions of these 3 fibril components over 24 hours. **(E)** STEM mass mapping electron microscopy data from mechanically dispersed entire Achilles tendons (n=4) showing the variation of the D1 fibril component at 3-hour time intervals during 24 hours. This is in the form of a double plot with the data repeated over 48 hours. The data were fitted by least squares minimization to a sinusoidal variation (red line). The grey panels indicate the night-time (dark) periods. **(F)** Still image from a step-through movie generated by serial block face-scanning electron microscopy of 6-week old mouse Achilles tendon harvested at CT3 (Supplementary Video 1). **(G)** Still from a step-through 3D reconstruction displaying individual fibrils in a small region of mouse tail tendon, showing merging of D1 fibrils onto larger D2 and D3 fibrils (Supplementary Video 2). Fibrils that appear in at least 80% of frames are colored. Images generated from serial block face-scanning electron microscope stack of transverse TEM images. Area shown is 4 µm by 4 µm, each frame is a longitudinal step of 100 nm through the tendon. **(H)** Selected images from a serial block face-scanning electron microscope step-through movie stack with a D3 fibril (outlined) surrounded by a D2 (dark blue) fibril and nine D1 fibrils (shown in different colors). Cut numbers are shown. **(I)** Circularity of collagen fibrils examined in transverse TEM images of Achilles tendons from ZT3 and ZT15 indicate fibril cross sections from ZT3 are more irregular than fibril cross sections from ZT15 (>900 fibrils measured in both ZT3 and ZT15 samples; ***p<0.001). Error bars show SEM.

### Rhythmic protein secretory pathway

We considered that the diurnal changes in collagen fibril diameters, fibril circularity, and tissue mechanical properties could be due to rhythmic changes in transcription, translation and secretion leading to rhythmic changes in ECM composition. To identify a molecular mechanism, we first examined data from time-series transcriptomic profiling of mouse tendon ^29^. The data showed that the expression of *Col1a1* and *Col1a2* mRNAs is not rhythmic in mouse tendon during 24 hours (Figure S2). Therefore, the diurnal behavior of collagen fibrils could not be explained by rhythmicity of collagen gene transcription. However, previous studies in liver had revealed circadian clock regulation of ribosome binding of the *Col1a1* transcripts ^32^. Here, we showed by electron microscopy that the packing of ribosomes along the ER was time-of-day dependent in mouse tendon fibroblasts *in vivo* and maximal at ZT15 (Figure 3SA, B). This was an indicator that circadian clock control of collagen homeostasis begins early in the secretory pathway by temporally regulating protein translation.

**Figure 3.**
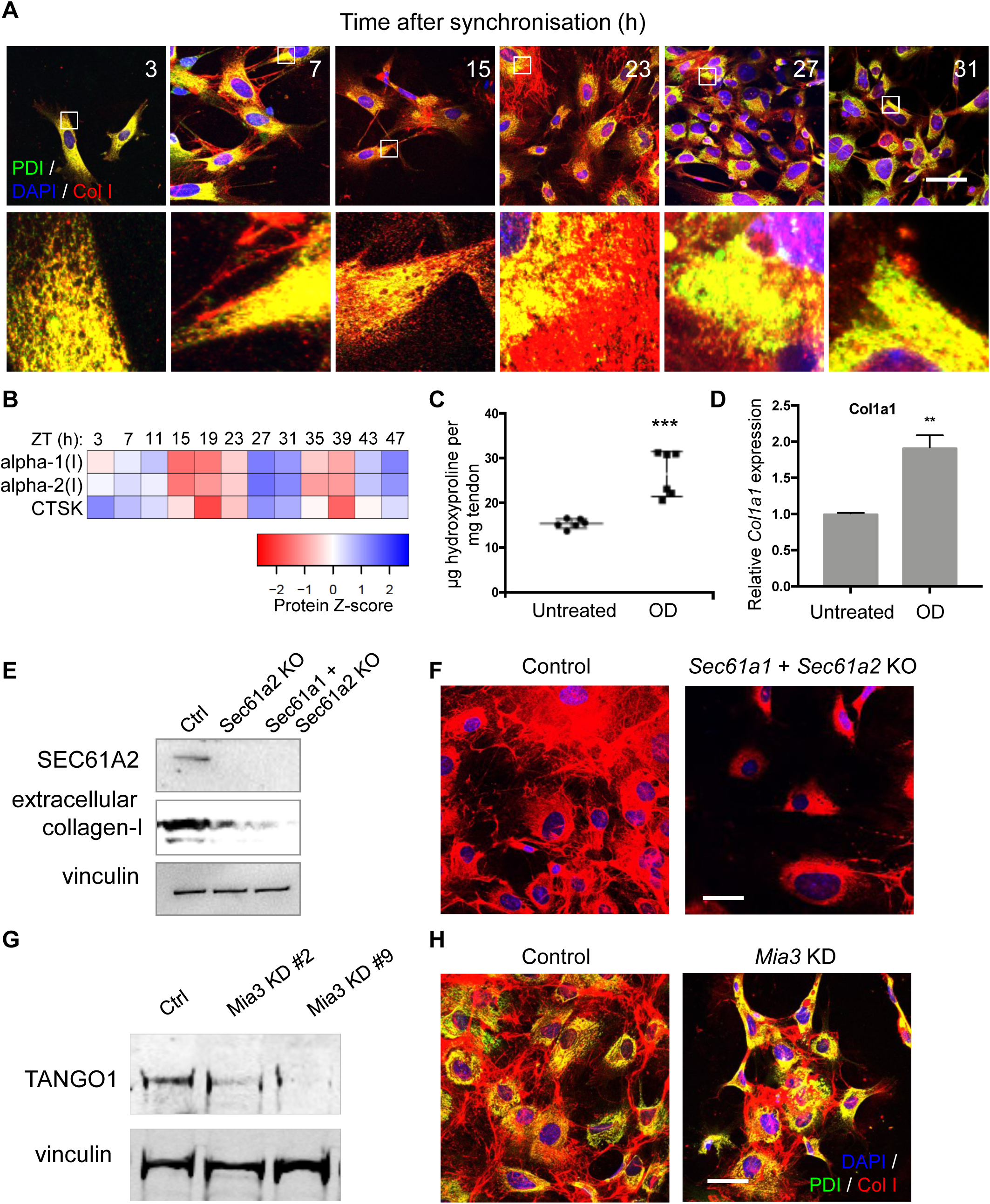
Circadian clock-regulated PC-I trafficking and collagen fiber assembly in culture. **(A)** Time series immunofluorescence study using anti-collagen-I and anti-PDI-specific antibodies. Areas boxed in white are magnified in the lower panels. Representative images from each time point are shown. **(B)** Heat map from LC-tandem mass spectrometry analyses of protein expression showing CTSK and collagen α1(I) and collagen α2(I) oscillation over 48 hours. **(C)** Increased hydroxyproline content of dissected *ex vivo* Achilles tendons from mice incubated with 1 µM odanacatib (OD) for 3 days compared to control (n=6; ***p<0.001). **(D)** *Col1a1* mRNA expression in mouse tail tendon fibroblast incubated with OD. **(E)** Western blot analyses of conditioned media from CRISPR/Cas9 *Sec61a2*-knock out and *Sec61a1/Sec61a2-*double knock out MEFs (*Sec61a2* KO, *Sec61a1* + *Sec61a2* KO) show that depletion of SEC61A2 inhibits collagen-I secretion. Levels of vinculin protein served as a loading control. **(F)** Immunofluorescence analysis of synchronized MEFs show that depletion of *Sec61a1* and *Sec61a2* prevent collagen-I fiber assembly in the extracellular matrix. **(G)** Western blot analysis of TANGO1 protein in MEFs treated with siRNA targeting *Mia3*. Levels of vinculin protein served as a loading control. **(H)** Immunofluorescence analysis of control and siMia3 cells shows collagen-I retention in the ER (PDI-staining) in cells with *Mia3* knock-down (*Mia3* KD). Staining of control cells show collagen-I fibers in the extracellular matrix. Bars 10 µm.

Intriguingly, transcripts encoding several key secretory pathway proteins were conspicuously rhythmic (Figure 2A). These included: *Sec61a2* (a core member of the translocon ^33^), *Mia3* (which encodes TANGO1 with a proposed role in facilitating cargo loading at ER exit sites ^34^), *Pde4d* (functionally related to another phosphodiesterase, *Pde7a1*, which has been proposed to be important in regulating PC-I transport in COS7 cells in culture ^35^), and *Vps33b* (a post-Golgi protein reported to be important in PC-IV post-translational modification ^36^). We also noted that the transcript encoding *Ctsk* (a collagenase important in collagen degradation ^37,38^) is 24-hour rhythmic. Western blot analysis showed that proteins encoded by these transcripts were rhythmic in adult mouse tendon (Figure 2B).

**Figure 2.**
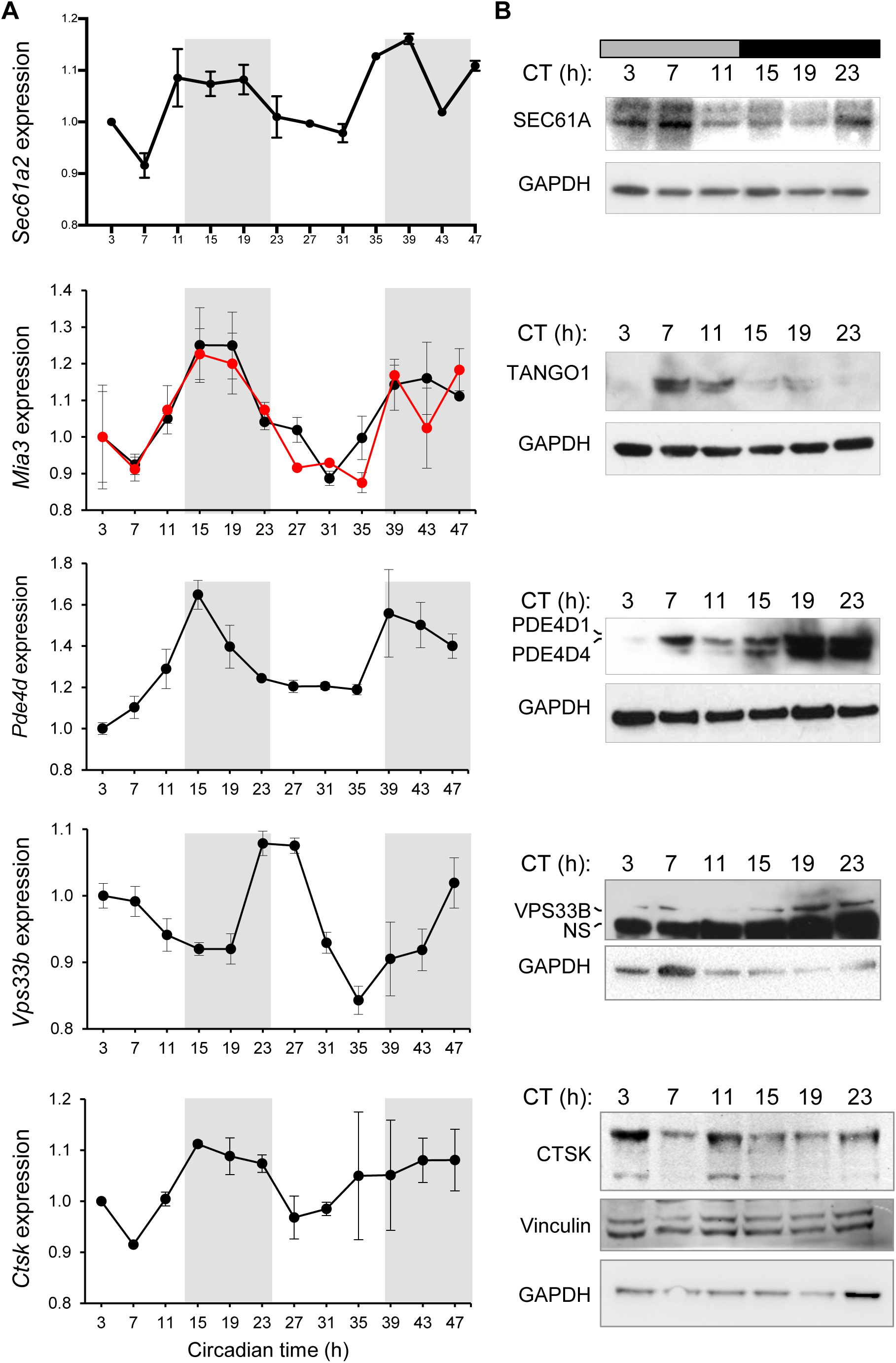
Rhythmic expression of genes regulating collagen trafficking and fiber assembly. **(A)** Rhythmic expression of *Sec61a2, Mia3, Pde4d, Vps33b* and *Ctsk* mRNA and (**B**) the corresponding proteins. Transcripts were detected by microarray over a 48-h period (multiple probe sets are shown in black and red, if present; n=2; bars show SEM). Protein was assessed over a 24-h period. Levels of GAPDH or vinculin served as loading controls. CT, circadian time. NS, non-specific staining.

### Circadian clock-synchronized cells in culture exhibit a rhythmic collagen matrix

Collagen secreting fibroblasts have been widely used as a model of peripheral clocks ^39^. Therefore, we used clock-synchronized cultured fibroblasts as an *in vitro* model system to evaluate the importance of *Sec61a2, Tango1, Pde4d, Vps33b* and *Ctsk* in collagen secretion and homeostasis. Using collagen I- and PDI-(ER-resident protein disulfide isomerase) specific antibodies, we showed that PC-I localizes to the ER early after synchronization (3 hours) and collagen fibrils begin to appear 7-15 hours later and are maximal at the end of the 24 hours post-synchronization (Figure 3A). The rhythmicity of collagen fibril assembly and disassembly continues for at least 3 days in culture (Figure S4). In further experiments, co-immunofluorescence studies using collagen I- and GM130-(an ERGIC/cis-Golgi resident protein) specific antibodies, showed a time-dependent localization of PC-I in ERGIC/cis-Golgi compartments (Figure S5). These experiments showed that a rhythmic secretory pathway was cell-intrinsic. Disappearance of some extracellular collagen fibrils at the end of the circadian cycle was indicative of a clock-regulated degradation step. Cathepsins are a family of cysteine proteases of which cathepsin K (CTSK) deficiency in pycnodysostosis results in accumulation of non-digested phagocytosed collagen in fibroblasts ^40^. *Ctsk* is transcriptionally and translationally rhythmic in tendon fibroblasts (Figure 2A, B and Figure 3B). Incubation of *ex vivo* tendons with odanacatib to inhibit CTSK resulted in collagen accumulation, as shown by quantitative hydroxyproline analysis (Figure 3C). Quantitative-PCR showed that odanacatib also led to increased expression of *Col1a1* mRNA (Figure 3D) suggestive of a feedback mechanism between collagen turnover and transcription.

### SEC61, TANGO1, PDE4D, and VPS33B are required for collagen fiber assembly

SEC61, TANGO1, PDE4D, and VPS33B are sequentially expressed during a circadian period and serially located within the secretory pathway in the direction of travel of secreted proteins. Thus, we postulated that they may have an essential function in regulating procollagen trafficking and collagen fiber assembly. To test this notion, we used knockdown and knockout strategies combined with immunolocalization fluorescence confocal microscopy.

*SEC61* The mRNAs of *Col1a1* and *Col1a2* (which encode α1 and α2 of type I collagen, respectively) have a 5’ UTR consisting of 2 stems flanked by a bulge (termed the collagen 5’ stem-loop) that binds LARP6 (La ribonucleoprotein domain family member 6) and are specifically targeted to the SEC61 translocon ^41^. The protein-conduction pore of the translocon is formed from SEC61α which associates with a β and a γ subunit to form a heterotrimeric complex SEC61αβγ in eukaryotes ^42–44 45^. There are two *Sec61a* genes (*Sec61a1* and *Sec61a2*), which encode SEC61α. CRISPR/Cas9-mediated knockout of *Sec61a1* and *Sec61a2* in MEFs (Figure 3E, Figure S3C) resulted in markedly fewer collagen fibers (Figure 3F). In the absence of the SEC61 translocon, transcription from *Col1a1* and *Col1a2* would be expected to continue, which raised the question about the fate of the proα1(I) and proα2(I) polypeptide chains. Co-immunofluorescence using anti-collagen-I- and anti-calnexin-specific antibodies showed that collagen was translated into the cytosol in the absence of the SEC61 translocon (Figure S3D).

*TANGO1* TANGO1 has been suggested to corral COPII coats to accommodate PC-I ^46^. We showed that siRNA-mediated knockdown of *Mia3*/TANGO1 in MEFs (Figure 3G and Figure S3E) resulted in reduced numbers of collagen fibers (Figure 3H). Knockdown of *Mia3*/TANGO1 resulted in co-localization of PDI (ER-resident protein disulfide isomerase) and collagen-I in the absence of extracellular collagen fibrils (Figure 3H).

*PDE4D* Previous studies in COS7 cells had shown that PDE7A1 regulates PKA-dependent retrograde transport of the KDELR in COS7 cells ^35^. HSP47 is an ER-resident molecular chaperone that contains a C-terminal KDEL signal and is required for procollagen secretion ^47^. Thus, we investigated if PDE4D regulated HSP47 transport. The results showed that HSP47 co-localized with GM130 (a marker of cis-Golgi/ERGIC) between 15- and 23-hours post-synchronization (Figure 4A). RNAi was highly effective at reducing the levels of *Pde4d* mRNA in MEFs (Figure S3F, G). Immunofluorescence using a collagen-I-specific antibody showed that *Pde4d* knockdown severely reduced the ability of cells to assemble collagen fibers and caused intracellular retention of PC-I, when compared to control cells (Figure 4B). In *Pde4d* KD cells, HSP47 showed poor co-localization with GM130 (Figure 4C), indicative of a blockage of HSP47 transport.

**Figure 4.**
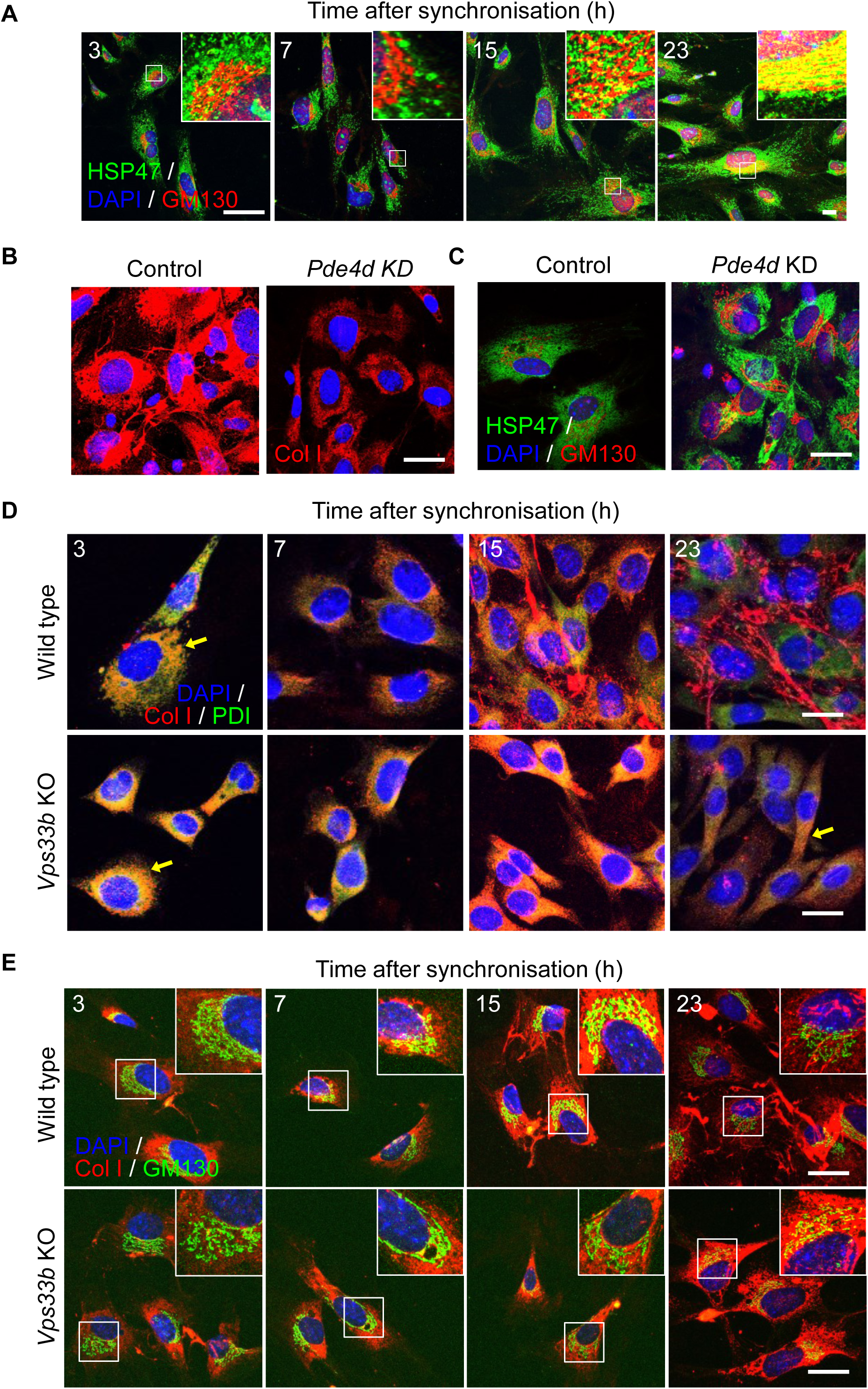
Golgi and post-Golgi transport of PC-I is rhythmic. **(A)** Immunofluorescence analysis of collagen chaperone HSP47, and ERGIC and the cis-Golgi (detected by marker GM130) in synchronized MEFs showed gradual co-localization of HSP47 with GM130 (see insets). **(B)** Immunofluorescence analysis shows that PDE4D depletion in MEFs using siRNAs (*Pde4d* KD) blocked collagen-I secretion into the matrix, which was unaffected in control cells. **(C)** Immunofluorescence analysis of HSP47 in *Pde4d* KD cells show accumulation of HSP47 in GM130 compared to control MEFs. **(D)** Immunofluorescence analysis of synchronized iTTFs shows collagen-I secretion (white arrows) was inhibited in CRISPR-mediated *Vps33b* KO cells, where collagen-I was retained in the ER (PDI staining) (yellow arrows). **(E)** Immunofluorescence analysis of synchronized MEFs also increased retention of collagen-I occurs in the Golgi (GM130 staining) when *Vps33b* was knocked out (see insets). Bars 10 µm.

*VPS33B* Post-Golgi trafficking of PC-I is poorly understood but recent studies suggest a role for VIPAR in the regulation of lysyl hydroxylation of collagen IV by PLOD3 (procollagen-lysine,2-oxoglutarate 5-dioxygenase 3) ^36^. The transcript and protein for *Vps33b*, a binding partner of VIPAR peak at the end of the circadian cycle (Figure 2A, B). Thus, we used CRISPR-Cas9 to knockout *Vps33b* in MEFs (Figure S3H) and immortalized tendon fibroblasts (iTTFs) (Figure S3H-J), and used collagen-I-, PDI- and GM130-specific antibodies to observe transport of PC-I through the secretory pathway during the circadian cycle. In non-CRISPR treated cells, PC-I co-localized with PDI 3 hours after synchronization (Figure 4D), with GM130 7-15 hours after synchronization (Figure 4E), and assembled collagen fibrils between 15- and 23-hours post-synchronization (Figure 4D, E). In contrast, PC-I co-localized with PDI throughout the circadian cycle, Also, whereas PC-I had transited through the Golgi by 23-hours post-synchronization in control cells, PC-I co-localized with GM130 at 23-hours post-synchronization in VPS33B knockout cells. Furthermore, numbers of collagen fibrils were severely reduced in VPS33B knockout cells (Figure 4D, E).

### A predictive mathematical model of the collagen secretory pathway

We used the above protein data, together with published data on mRNA transcriptional rates ^48^, to generate a predictive model of the transport of PC-I through the secretory pathway. We built a deterministic ordinary differential equation system based on the principles of mass-action enzyme kinetics, with variables representing cellular concentrations of collagen and relevant proteins in ER (translation via SEC61 with phase of CT23-7), ER exit sites (pathway progression controlled by TANGO1 and HSP47, CT7-11), Golgi (PDE4D directed transport of HSP47 at CT15-23), post-Golgi (pathway progression controlled by VPS33B, CT19-23), and plasma membrane (release step at CT0-3). Collagen degradation is modeled based on cathepsin K (CTSK, CT15-27). Simulations and subsequent analysis were carried out using the differential equation solver *NDSolve* in Wolfram Mathematica 11.3 (see STAR★METHODS). The simulation predicted that the relative amounts of PC-I in ER, ER exit sites, Golgi, post-Golgi compartments and the extracellular matrix varied depending on the circadian cycle (Figure 5A and Video 5). The model predicted low levels of PC-I throughout the circadian cycle in ER exit sites (compared to other transport compartments) (Supplementary Video 5), which was due to the narrow window of time when TANGO1 was expressed in the circadian period and the rapidity of microtubule transport of vesicles. The model predicted an inverse relationship (to a close approximation) between intracellular PC-I and extracellular collagen concentration (Figure 5B), assuming that PC-I is converted to collagen-I during transport or in the ECM, which agreed with previous observations ^49,50^. However, the shape of the phase curve (Figure 5B) predicted a 6-hour time delay between the peak in PC-I and the peak in collagen-I. Importantly, the model predicted that extracellular collagen-I was rhythmic within a circadian period, with the peak of collagen occurring in the presumptive day (Figure 5C).

**Figure 5.**
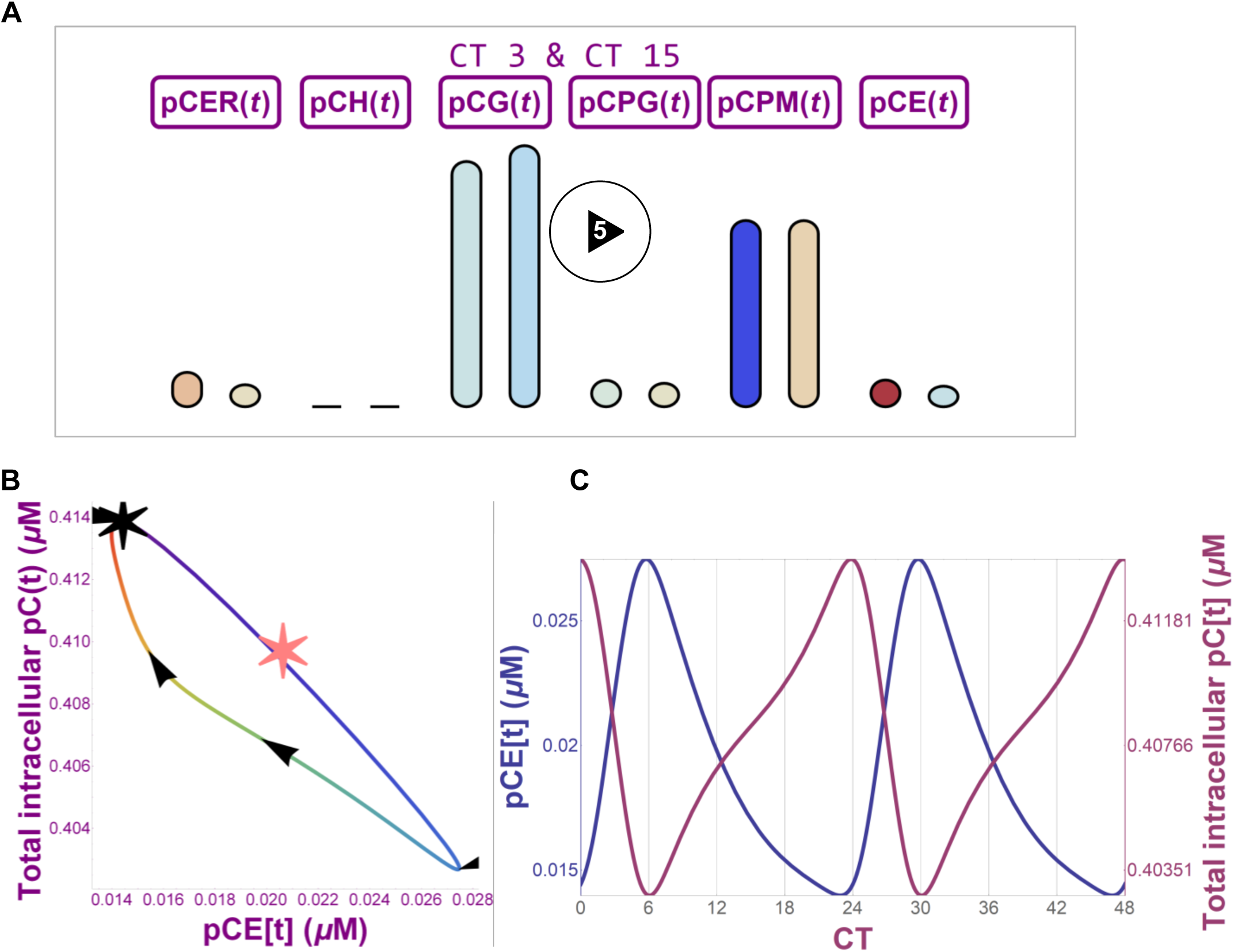
Mathematical prediction of PC-I and collagen I secretion. **(A)** Still frame from a video representation (Supplementary Video 5) of a mathematical model of the secretory pathway showing PC-I levels in ER (SEC61-dependent translation of PC-I into ER, pCER(t)), ER exit sites (TANGO1-dependent PC-I export from ER, pCH(t)), ERGIC (PDE4D-dependent transport, pCG(t)), post-Golgi (VPS33B-dependent transport, pCPM(t)) and extracellular matrix (CSTK-dependent collagen degradation, pCE(t)). Bar height represents collagen concentration, and color represents the rate of change of concentration (blue - low to red - high). The paired bars represent collagen levels at CT3 and CT15. **(B)** Phase portrait showing relationship between intracellular collagen concentration and extracellular collagen concentration over a 24-hour cycle. Black star shows CT0, and the line color shows the circadian time increasing in the direction indicated by the arrows. The pink star indicates stationary concentration values for the model when the circadian forcing functions are removed. **(C)** 48-hour time plot (CT0 to CT48) of intracellular (red) and extracellular (blue) collagen concentration fluctuations as predicted by mathematical modeling.

### Rhythmic proteome revealed daily surge of collagen production in tendons *in vivo*

To explore the prediction that a rhythmic secretory pathway generates a wave of collagen-I in the day, we performed a time-series proteomic analysis of mouse tendon using an extraction buffer that solubilized non-covalently bound collagen whilst leaving the permanent, cross-linked collagen intact (see STAR**★**METHODS). This allowed for separate analysis of newly-synthesized collagen from the permanent collagen. Electron microscopy showed the persistence of D2 and D3 fibrils, and depletion of D1 fibrils, in the non-solubilized (pellet) fraction (Figure S6A-C). Hydroxyproline analysis showed that the solubilized collagen accounted for 18% of total collagen (Figure S6D). The soluble proteins were treated with trypsin and the resultant peptides analyzed by liquid chromatography-tandem mass spectrometry ^51^. We detected ∼10,000 peptides and identified 1,474 proteins with ≥ 2 peptides per protein (Figure 6A) (Supplementary Data 1), with good reproducibility between biological replicates as indicated by Pearson analysis across two consecutive 24-hour periods (Figure 6B). Rhythmic profiles of >100 proteins were detected during 48 hours (Figure 6C), with prominent changes occurring at the night-day and day-night transitions for rhythmic and non-rhythmic proteins (Figure 6D, E and Figure S7A). Protein ontology analysis showed that actin-binding proteins were most abundant in night samples (Figure S7B). This supports a recent proteomics study in fibroblasts, which indicated regulation of actin dynamics is under circadian control ^52^. Approximately 10% of detected proteins (141) were robustly rhythmic (with periods of 20-27.5 h) (Figure 6F). The α1(I) and α2(I) chains of collagen-I were conspicuously rhythmic and peaked during daylight hours, equivalent to 1.6-fold difference in extractable collagen between day and night sampling (Figure 6G). It was possible to identify peptides specific for the C-propeptide region of the proα1(I) proα1(I) chain of PC-I, which peaked ∼6 hours prior to the α1(I) and α2(I) peptides and was consistent with the slow cleavage of PC-I to collagen-I ^53,54^ (Figure 6H and Supplementary Data 2). To investigate the rate of collagen-I production between the day and night, tendon tissues dissected 12 hours apart were labeled with ^14^C-proline *ex vivo* ^30^. Both Achilles and tail tendon tissues had significantly greater PC-I production at night (Figure 6I and Figure S7C, D), corroborating the LC-tandem mass spectrometry data. ^14^C-amino acid labeling of newly-synthesized proteins every 4 hours during 24 hours confirmed that PC-I production was ramped up throughout the night (Figure S7E, F).

**Figure 6.**
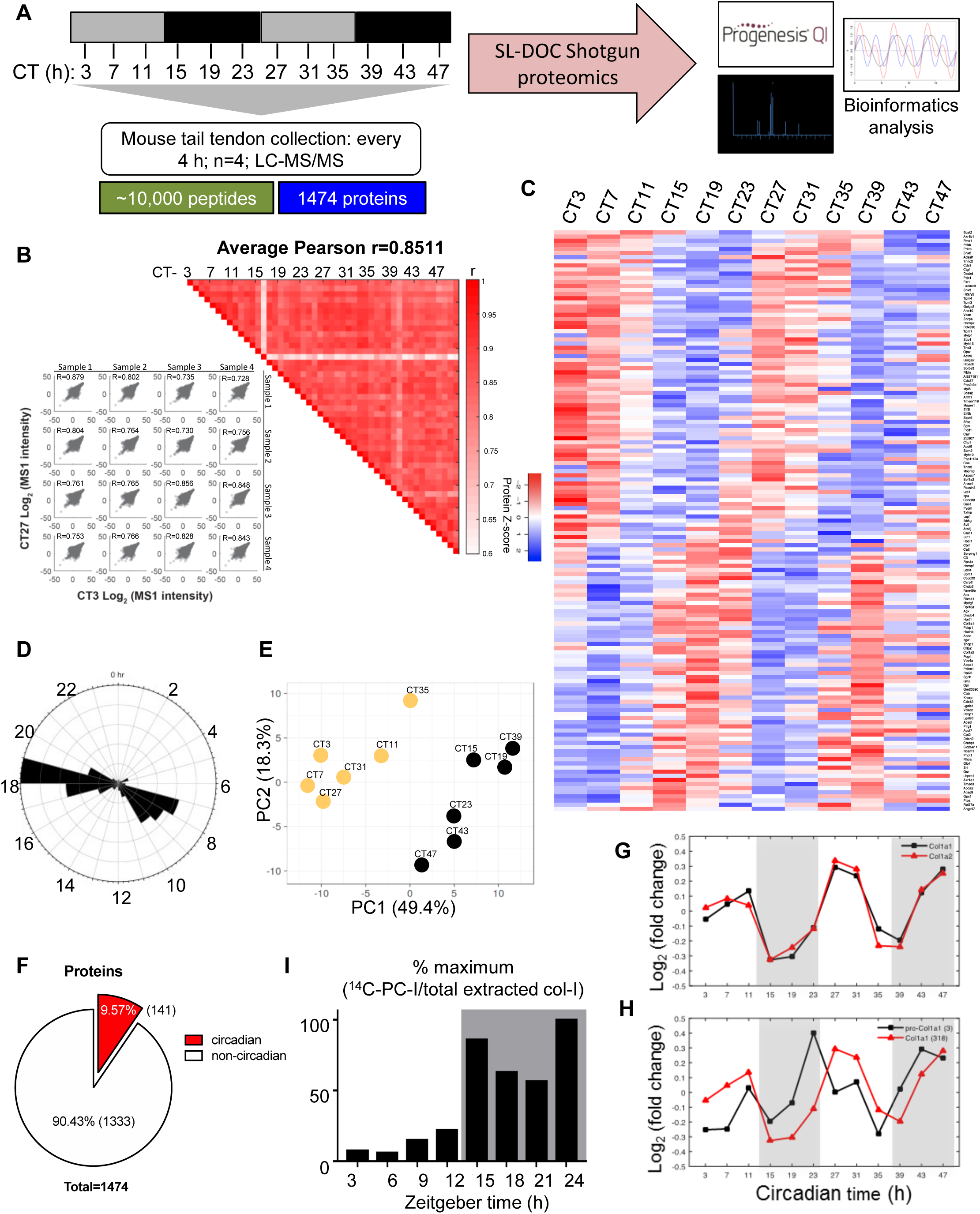
Time-series proteomics analysis of proteins in mouse tendon. **(A)** Schematic showing collection of mouse tail tendons every 4 h during 48 h for proteomics analysis. **(B)** Heat map of Pearson correlation coefficients (r) for pairwise comparison of each sample, calculated from normalized peptide intensities. Each square represents a single comparison with intensity of color denoting r according to the color bar (right). High average and minimum r values denote high reproducibility between biological replicates (left). **(C)** Heat map depicting the levels of the detected rhythmic proteins, organized according to the timing of peak expression in circadian time (CT). Red is downregulated and blue is up-regulated based on protein Z-score. **(D)** Radial plot showing the frequency distribution of the phases of the rhythmic proteome in mouse tendons. The majority of peptides were detected between CT6–9 and CT17–19. **(E)** Principal component analysis (PCA) of the cycling proteome. The major two components separating the data organized the samples into samples corresponding to day (left, yellow circles) and night (right, black circles) time points. **(F)** Pie chart indicating the percentage of circadian proteins and non-circadian proteins. **(G)** Relative abundance of the two collagen-I α chains, α1 (Col1a1) and α2 (Col1a2), detected by mass spectrometry over 48 h (mean of log10 intensities for n=4). Peak abundance of collagen-I peptides was detected at CT27 and CT47. **(H)** Peptide counts (log_2_ values) of C-propeptide region of α1 (pro-Col1a1) chain and of α1 (Col1a1) detected by mass spectrometry over 48 h, based on calculations using peptides from the different sub-domains of pro-α1(I) chain. Note the abundance of C-pro-α(I) peptides peaks prior to the peak abundance of α1(I) peptides, at CT23 and CT43. **(I)** Achilles and tendon tissues were harvested at the day phase (zeitgeber time (ZT) 3, day) or night phase (ZT15, night) and labeled with ^14^C proline for 1 h in the presence of exogenous L-ascorbate. Densitometry of ^14^C-labeled collagen-I from western blot analysis showed significant increase in ^14^C-labeled collagen-I in the night phase.

### Mistiming of the secretory pathway results in collagen accumulation and abnormal fibril structure

If a 24-hour rhythmic secretory pathway generates a daily wave of collagen and rhythmic assembly of collagen fibers (as shown in Figure 3A), an arrhythmic pathway may be expected to result in arrhythmic or non-24-hour cyclical synthesis of collagen fibers. To test this prediction we cultured fibroblasts from wild type and *ClockΔ*19 mice bred on a *PER2::LUC* background, and treated the cells with the clock synchronizing agent dexamethasone. *ClockΔ19* mice have disrupted circadian rhythms ^55^ and the in-frame fusion of luciferase at the *Per2* locus enabled direct measurement of the endogenous circadian rhythm. Wild type cells responded well to the synchronizing effect of dexamethasone and showed a robust bioluminescence rhythmicity that persisted for 3 days (Figure 7A). In contrast, *ClockΔ19* fibroblasts did not sustain a rhythm (Figure 7A). Time-series quantitative-PCR analysis of *Nr1dr1* confirmed that dexamethasone was effective in synchronizing the clock in wild type cells only but not in *ClockΔ19* cells (Figure 7B). Levels of *Col1a1* mRNA were not rhythmic in synchronized wild type *ClockΔ19* cells (data not shown), which agreed with *in vivo* data (see Figure S2). However, the overall levels of *Col1a1* mRNA were elevated (Figure 7C) and the overall levels of *Ctsk* mRNA were decreased (Figure 7D) in *ClockΔ19* compared to wild type cells.

**Figure 7.**
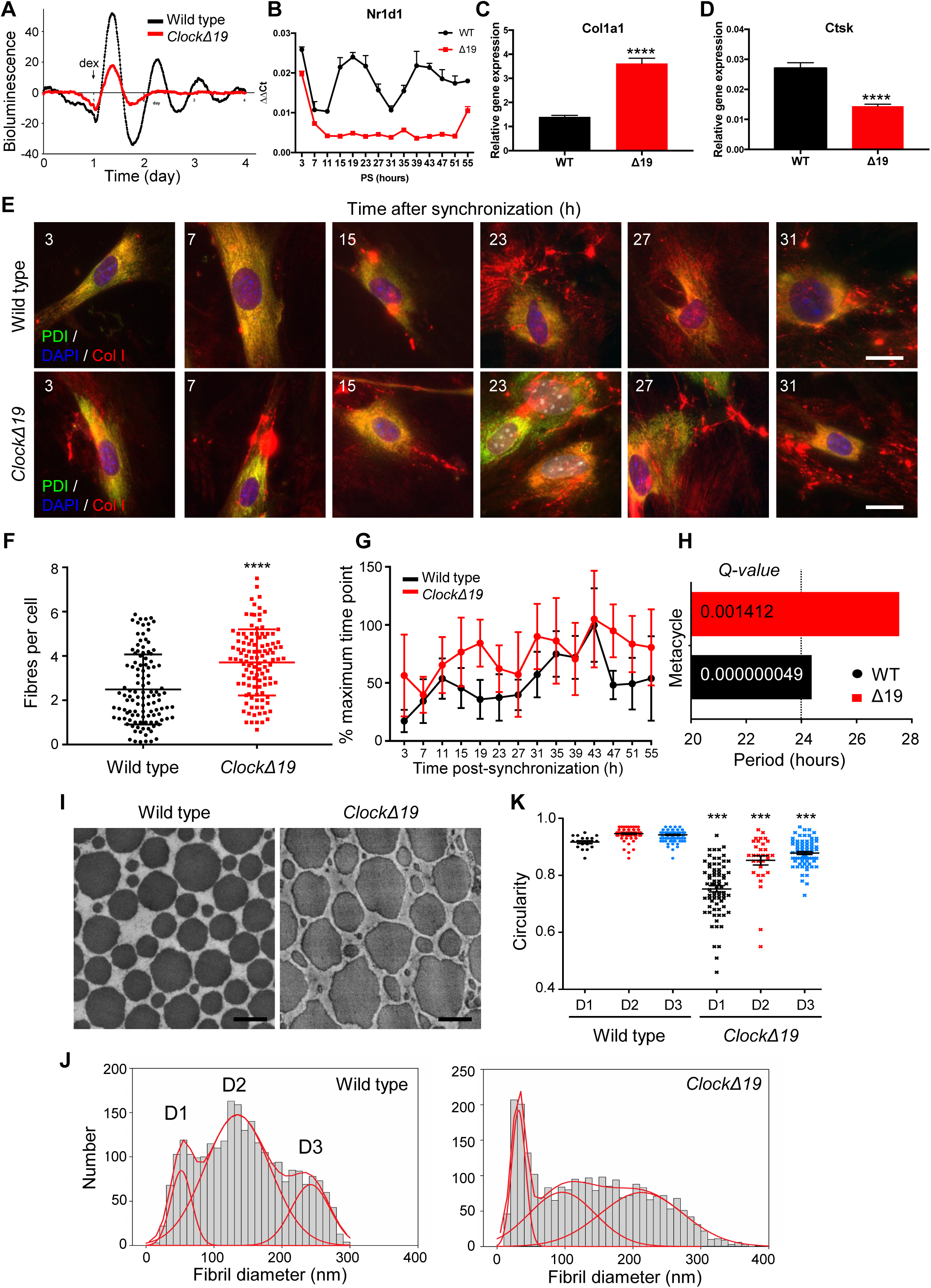
Dysregulated collagen fiber homeostasis in the absence of a circadian rhythm. (**A**) Luminometry recordings of endogenous circadian rhythms and re-initiation of the rhythms with dexamethasone (arrow) evident in extracted fibroblasts from wild type, but not *ClockΔ19* mice bred on a *PER2*::Luciferase background. **(B-D)** Gene expression analysis **(B)** *Nr1d1* **(C)** *Col1a1* (average of all time points) **(D)** *Ctsk* (average of all time points) in fibroblasts obtained from wild type and *ClockΔ19* mice. RNA was collected every 4 h over 55 h after synchronization with dexamethasone. **(E)** Immunofluorescence analysis showed more and split collagen-I fibril assembled by the fibroblasts of *ClockΔ19* mice. Bars 10 µm. **(F)** Total number of fibrils per cell measured, indicating that *ClockΔ19* fibroblasts produced more fibrils overall. **(G)** Plot showing percentage collagen fiber count in fibroblasts isolated from wild type and *ClockΔ19* during 55 hours, corrected to the maximum percentage fibril count of the time course. *ClockΔ19* fibroblasts have lost their rhythmicity in fibril assembly. **(H)** MetaCycle analyses of fibril counts showed a *circa* 24-h rhythmicity in wild type fibroblasts, compared to *circa* 28-h rhythmicity in *ClockΔ19* fibroblasts. **(I)** Representative transverse transmission electron microscopy images of tail tendons from 6 weeks-old wild type and *ClockΔ19* showing irregular fibril cross sections in mutant tendons. Bars 200 nm. **(J)** Circularity of D1, D2 and D3 collagen fibrils of tail tendons sampled from *ClockΔ19* and their wild type littermate controls (192 wild type and 179 *ClockΔ19* fibers measured. ***p<0.0001 in D1, D2 and D3 populations in mutants compared to same populations in littermate controls). **(K)** Fibril diameter distributions measured from transverse TEM images of wild type and *ClockΔ19* tail tendons showing abnormal distribution of fibril diameters and increase in large (>400-nm fibrils) in mutant tendons (>1000 fibrils measured). Red lines show 3-Gaussian fit curves.

Next, we synchronized wild type and *ClockΔ19* fibroblasts and performed a time-series immunofluorescence study to quantitate the number of collagen fibers formed (Figure 7E). The results showed that wild type and *ClockΔ19* fibroblasts synthesized collagen fibers during 55 hours in culture; therefore, a rhythmic secretory pathway is not necessary for collagen fiber formation. However, *ClockΔ19* fibroblast cultures accumulated more fibers than wild-type cells (Figure 7F). Metacycle analysis of the numbers of fibers indicated that the increase and decrease in fiber number during days in culture in wild type and *ClockΔ19* cultures was 24 and 28 hours, respectively (Figure 7G).

the number of collagen fibers synthesized by wild-type cells increased and decreased to reach a steady-state level after 55 hours (Figure 7F). In contrast, the number of collagen fibrils synthesized by *ClockΔ19* fibroblasts increased and decreased with a periodicity of 28 hours (Figure 7G) and increased during 55 hours without reaching a plateau (Figure 7F). As a consequence, the number of collagen fibers per cell was greater in *ClockΔ19* than in wild-type fibroblasts (Figure 7H).

The *ClockΔ19* is a global mutant mouse that shows systemic loss of rhythmicity, with potential confounding factors. To selectively disrupt circadian rhythm in tendon-lineage cells we generated a new line of *Scx-Cre::Bmal1 lox/lox* knockout mice that lacked circadian rhythm in tail and Achilles tendons (Figure S8A, Videos 6 (wild-type) and 7 (*Scx-Cre::Bmal1 lox/lox*)). Using TEM, we showed that the tendon fibrils in *ClockΔ19* and *Scx-Cre::Bmal1 lox/lox* mice were irregular in outline (Figure 7I and Figure S8B), had disrupted fibril diameter distributions (Figure 7J and Figure S8C) and decreased circularity values (Figure 7K and Figure S8D). Tendons in clock-disrupted mice were significantly thicker and had reduced elastic modulus and maximum load compared to wild type littermates (Figure S8E and F). There were no differences in fibril volume fraction between arrhythmic and wild type tendons (Figure 8SE); hence, the increased tendon width in the mutant mice was due to an increased volume of collagen. These results showed that disrupting the circadian clock results in increased collagen deposition, decreased elastic modulus, decreased maximum load, and irregular fibril structure, which demonstrate disrupted matrix homeostasis.

## Discussion

In our study we have shown a rhythmic pool of collagen in the extracellular that results from circadian clock regulation of the protein secretory pathway. Mistiming of the pathway in cells and mice lacking a functional circadian clock resulted in dysregulated collagen homeostasis as seen in fibrosis. The synthesis and removal of a distinct pool of collagen whilst leaving the bulk protein unchanged illustrates a surprisingly dynamic matrix and a selective mechanism of protein homeostasis that maintains tissue function and organization.

Intracellular proteins are constantly being synthesized and degraded in a cyclical process arising from transcription, translation and protein folding to replace old and damaged polypeptides with newly-synthesized copies ^56^. However, the extreme longevity of collagen in tissues would suggest that these conventional transcriptional and translational homeostatic mechanisms do not apply to collagen. Presumably, the energy- and time-dependent developmental processes that are required to establish extensive networks of collagen fibrils in tissues (which in humans equates to ∼17 years) rule out a protein homeostatic mechanism in which all of the collagen is turned over. Tissues that are rich in collagen are some of the highest mechanically-loaded tissue in vertebrates and are exposed to repeated cycles of stress and strain throughout the life of the animal. Therefore, collagen molecules that are never replaced would be expected to undergo mechanical damage leading to fatigue failure. A mechanism of protein homeostasis must therefore exist to protect, rather than replace, the collagen that was assembled during skeletal growth. Such a mechanism must be able to account for the observed continued synthesis of collagen in the face of zero turnover of the bulk of the tissue collagen. The circadian clock mechanism described here would appear to fulfill these requirements.

Our study showed that a population of thin (D1) fibrils are interspersed between thick (D2 and D3) collagen fibrils, and make frequent contact with the surfaces of the thick fibrils. It has been suggested that the thin fibrils are part of a sheer stress loading mechanism in tendon ^57^. Our data would agree with this suggestion, particularly as the D1 fibrils do not appear to be under tensile strain. However, the fact that the proportion of D1 fibrils varies with time of day, and that the collagen-I in these fibrils is circadian clock rhythmic, may suggest an additional function to maintain matrix homeostasis in the face of bouts of activity and long-term wear and tear on the tissue. Collagen fibrils are ‘molecular alloys’ that comprise a variety of collagen types ^58^ and associated molecules including small leucine rich proteoglycans (SLRPs) (for a review of SLRPs, see ^59^). We noted that collagen α2(V) was rhythmic in-phase with collagen α1(I) and collagen α2(I) (Supplementary Data 1). Collagen-V controls the initiation of collagen fibril assembly, and haplo-insufficient mice have fewer, large and irregular collagen fibrils ^60^. In addition, decorin (an abundant SLRP found in a wide range of tissues) was rhythmic, peaking at CT21 (Supplementary Data 1). Mice deficient in decorin have weakened tissues and abnormal collagen fibrils with irregular profiles ^61^. This raises the possibility that the rhythmic collagen-I-rich D1 fibrils are part of a multicomponent chronomatrix.

Early work using electron microscopy, histochemistry and autoradiography demonstrated that procollagen molecules could be present in the Golgi apparatus and post-Golgi secretory vesicles within 20-30 mins of synthesis in the ER ^62^. Later studies using ^14^C-labeled amino acids showed that PC-I could be synthesized and secreted within 20 mins ^30,50^. The need to use radiolabel is these studies to detect the procollagen suggests that the concentrations of procollagen were low, and perhaps below the critical concentration for collagen fibril assembly ^63^. Taken together, cells have the ability to secrete low levels of procollagen rapidly (∼minutes) and to increase the concentration of procollagen being delivered to the plasma membrane by circadian clock regulation of SEC61, TANGO1, PDE4D and VPS33B. Presumably, damming procollagen flow from one secretory compartment to the next is a mechanism of increasing procollagen concentration to facilitate D1 collagen fibril assembly. The existence of two temporal mechanisms of procollagen secretion might have implications for day time protection of the collagenous matrix following bouts of nocturnal activity in mice. Furthermore, the absence of rhythmic collagen homeostasis, or a shift to continuous collagen secretion, might be a disease mechanism in poor wound healing, fibrosis, and musculoskeletal degeneration.

## METHODS

### CONTACT FOR REAGENT AND RESOURCE SHARING

Further information and requests for reagents may be directed to, and will be fulfilled by the corresponding author Karl Kadler.

A separate resource document is uploaded.

## EXPERIMENTAL MODELS

### Mice

The care and use of all mice in this study was carried out in accordance with the UK Home Office regulations, UK Animals (Scientific Procedures) Act of 1986 under the Home Office Licence (#70/8858 or I045CA465). The permission included the generation of conditional knock out animals.

*ClockΔ19* mice (C57BL/6 on a PER2::Luciferase reporter background) were as described previously ^55^. To generate mice in which *Bmal1* is ablated in tendon-lineage cells, we crossed mice expressing Cre recombinase under the control of the Scleraxis promoter (*Scx-Cre*; C57BL/6) ^64^ with C57BL/6 mice carrying loxP sites flanking the exon encoding the BMAL1 basic helix-loop-helix domain ^65^ bred on a background with PER2::Luciferase reporter ^66^ to produce *Scx-Cre::Bmal1 lox/lox* mice. Six-week old wild type C57BL/6 mice housed in 12-h dark/12-h dark cycle (DD) were used for circadian time (CT) time-series microarray and time-series protein extraction for western blotting as previously described (Yeung et al., 2014). Wild type C57BL/6 mice housed in 12-h light/12-h dark cycle (LD) were used for all remainder zeitgeber time (ZT) studies, including time-dependent electron microscopy experiments, cyclic loading mechanical testing, *ex vivo* ^14^C-proline labelling experiments, proteomics analyses, and *ex vivo* odanacatib treatment.

## METHOD DETAILS

### Preparation of Cells

Tail tendon and lung fibroblasts from wild type and *Clock*Δ*19* were released from 6-8-week mouse tail tendons by 1000 U/ml bacterial collagenase type 4 (Worthington Biochemical Corporation) in trypsin (2.5 g/L) as previously described ^67^. Wild type tail tendon fibroblasts were immortalized by retroviral expression of mTERT (iTTFs) ^68^. Mouse embryonic fibroblasts (MEFs) were isolated from E14 wild type mouse embryos. Cells were cultured in complete media (Dulbecco’s modified Eagle’s medium: Nutrient Mixture F-12 containing 4500 mg/L glucose, L-glutamine, and sodium bicarbonate, supplemented with 10% fetal calf serum, 200 μM L-ascorbic acid 2-phosphate, 10,000 U/ml penicillin/Streptomycin) at 37°C, in 5% CO_2_.

To synchronize the circadian clocks of a population of cells, cells were incubated with either 100 nM dexamethasone (Merck) in complete media for 30 min at 37°C, in 5% CO_2_ or 50% horse serum (Thermo Fisher Scientific) in DMEM F-12 for 2 h at 37°C, in 5% CO_2_ and then for 24 h in complete medium at 37°C, in 5% CO_2_. After (at 0 h after synchronization), cells were fixed for immunofluorescence or used for protein or RNA isolation, at 3-h or 4-h intervals.

### CRISPR-Cas9-Mediated Knockout

Cells were treated with CRISPR-Cas9 to delete *Sec61a1, Sec61a2* or *Vps33b* genes. Ribonucleoprotein complexes of crRNA, tracrRNA and recombinant 3xNLS Cas9 were formed according to manufacturer’s protocol (Integrated DNA Technologies). Two different guides were used in combination to target *Sec61a1* (g1: TCC GGC AAG ATG ACA CAG AA, g2: TCT GCA AAA AGG GTA CGG CT), *Sec61a2* (g1: AGT TTA GAG AGA AGG TAC TA, g2: TTT ATG ATC GCT CAC ATT TC) or *Vps33b* (g1: CGT TAG CAA TTC GAT CCA AA, g2 AGT CAT CCT AAG CCC TCA GA). 60 pmol complexes (2 pmol Cas9) were nucleofected into 4 × 10^5^ MEFs using P1 Nucleofection Buffer (Lonza). Gene knockout was confirmed by western blotting and qPCR. For iTTF studies, 5 × 10^4^ cells were plated out in a 24-well plate, and the aforementioned ribonucleoprotein complexes were introduced into the culture using Lipofectamine RNAiMAX reagent (Thermo Fisher Scientific), according to manufacturer’s instructions. Clones were then generated using serial dilution in a 96-well plate. Gene knockdown was confirmed by western blotting and qPCR.

### RNA Interference

Cells were treated with siRNAs to deplete *Mia3* or *Pde4d* transcripts in mouse cells. siRNAs targeting Pde4d (SMARTpool ON-TARGETplus *Pde4d* siRNA; Dharmacon) or *Mia3* (GeneSolution *Mia3* siRNA; Qiagen) were transfected into MEFs using MEFs Nucleofector Kit (Lonza) according to the manufacturer’s instructions. Briefly, 4 × 10^6^ cells (in a well of a 6-well plate) were transfected with a 0.8 nmol of siRNA in a final volume of 3 mL (yielding a final concentration of 250 nM).

### RNA isolation and Quantitative Real-Time PCR

RNA was isolated from *ex vivo* tendon and cells as described previously ^67^. In brief, tissues and cells were disrupted in TRIzol Reagent (Thermo Fisher Scientific) and RNA was isolated according to manufacturer’s protocol. DNase treatment of RNA was performed using RQ1 RNase-Free DNase (Promega Corporation) according to manufacturer’s protocol. RNA integrity was assessed by gel electrophoresis and RNA concentration was measured using a NanoDrop 2000 (Thermo Fisher Scientific). Complementary DNA was synthesized from 2 µg RNA using TaqMan Reverse Transcription Reagents (Thermo Fisher Scientific).

SensiFAST SYBR kit reagents were used in qPCR reactions. Primer sequences used were *Col1a1* GCC TGC TTC GTG TAA ACT CC and TTG GTT TTT GGT CAC GTT CA, *Ctsk* GGA CCC ATC TCT GTG TCC AT and CCG AGC CAA GAG AGC ATA TC, *Gapdh* AAC TTT GGC ATT GTG GAA GG and ACA CAT TGG GGG TAG GAA CA, *Mia3* AGC CAC GGA CGG CGT TTC TC and CCC CTG CCA GTT TGT AGT A, *Nr1d1* CTT CCG TGA CCT TTC TCA GC and GGT AAG TGC CAG GGA GTT GA, *Pde4d* CCA ACT AGC CTC CCC TTG AA and GCT GTC AGA TCG GTA CAG GA, *Sec61a1* CAG GTT CTT CTG CCA AAG ATG and TCA TGC ACC ATG GAG GTC T, *Sec61a2* TGT ATG TCA TGA CGG GGA TG and CAA ACC AGC AAC AAA CAA CTG, and *Vps33b* GCA TTC ACA GAC ACG GCT AAG and ACA CCA CCA AGA TGA GGC G. Primers were validated by DNA sequencing of PCR products and verified using BLAST. For qPCR analysis, the 2^-ΔΔCt^ method ^69^ was used to analyze relative fold changes in gene expression compared to the first time point or to wild type or control.

### Protein Extraction and Western Blotting

Protein was extracted and analyzed by western blotting as previously described ^29^. Briefly, tendons were snap frozen and homogenized with ice-cold cell lysis buffer (20 mM Tris/HCl pH 7.6, 150 mM NaCl, 1 mM ethylene diamine tetra-acetic acid (EDTA), 1% (v/v) Igepal CA-630, 50 mM sodium fluoride) containing complete, mini EDTA-free protease inhibitor cocktail and PhosSTOP phosphatase inhibitor cocktail (Roche). Lysates were cleared by centrifugation at 10000 × g, at 4°C. Pierce BCA Protein Concentration Assay (Thermo Fisher Scientific) was used to determine protein concentration.

Proteins were separated on NuPAGE Novex 10% polyacrylamide Bis-Tris gels with 1x NuPAGE MOPS SDS buffer (both from Thermo Fisher Scientific) and transferred onto PVDF membranes (GE Healthcare). Membranes were blocked in 5% skimmed milk powder in 1x PBS containing 0.05% Tween 20. Antibodies were diluted in 2.5% skimmed milk powder in 1x PBS containing 0.05% Tween 20. Primary antibodies used were goat pAb to collagen α1 type I (1:1000; Santa Cruz), mouse mAb to GAPDH (1:10000; Merck), rabbit pAb to PDE4D (1:500; Santa Cruz), rabbit pAb to Sec61A (1:500; Santa Cruz), rabbit pAb to Tango1 (1:500; Santa Cruz), mouse mAb to vinculin (1:200; Oncogene), and mouse mAb to VPS33B (1:500; Proteintech). HRP-conjugated antibodies and Pierce ECL Western Blotting Substrate (both from Thermo Fisher Scientific) were used and reactivity was detected on Hyperfilm (GE Healthcare).

### Immunofluorescence and Laser Confocal Imaging

Cells were fixed with 100% methanol at −20°C and permeabilized with 0.5% Triton-X/PBS. Collagen-I was detected using a rabbit pAb (1:500; Gentaur), HSP47 was detected using mouse mAb (1:100; Santa Cruz), ER was detected using rabbit pAb to calnexin (1:200; Abcam) or mouse mAb protein disulfide isomerase (PDI) (1:100; Abcam), cis-Golgi was detected using rabbit mAb to GM130 (1:250; Abcam) or mouse pAb to GM130 (1:500; 610822, BD Biosciences). Secondary antibodies conjugated to FITC (Abcam) and Cy5 (Thermo Fisher Scientific) were used. Stained cells were mounted using VECTASHIELD Hardset antifade mounting medium with DAPI (Vector Labs).

Images were collected on a Leica TCS SP5 AOBS inverted confocal using an [63x / 0.50 Plan Fluotar] objective [and 3x confocal zoom]. The confocal settings were as follows, pinhole [1 airy unit], scan speed [1000 Hz unidirectional], format 1024 × 1024. Images were collected using [PMT] detectors with the following detection mirror settings; [FITC 494-530 nm; Cy5 640-690 nm] using the [488 nm (20%), 594 nm (100%) and 633 nm (100%)] laser lines respectively. To eliminate crosstalk between channels, the images were collected sequentially. When acquiring 3D optical stacks the confocal software was used to determine the optimal number of Z sections. Only the maximum intensity projections of these 3D stacks are shown in the results.

### *Ex Vivo* Tendon Culture

Following euthanization hind legs were amputated just below the knee. The skin was removed to expose the Achilles tendon. The whole bone-muscle-tendon unit (tibia, calcanium, gastrocnemius and soleus muscles with the Achilles tendon) was placed in complete media containing 1 µM odanacatib (Cambridge Bioscience Ltd.) or 0.1% DMSO vehicle control. *Ex vivo* tendons were cultured for 72 h at 37 °C, in 5% CO_2_.

### ^14^C-Proline Labelling of *Ex Vivo* Tendons

Achilles tendons were dissected from mice, rinsed in PBS, and incubated at 37°C in complete media containing 2.5 Ci/ml of ^14^C-proline (Perkin Elmer) for one hour, with the tubes agitated every 20 min. After incubation, tissues were washed, spun down and transferred to fresh Eppendorf tubes with 100 μl of salt extraction buffer (1 M NaCl, 25 mM EDTA, 50 mM Tris-HCl, pH7.4) containing protease inhibitors and phosphatase inhibitors. Tendons were extracted in 2 changes of salt extraction buffer – 12 h (S1), 6 h (S2) – these steps extracted collagen that was present in the extracellular fraction of the tissue. This was then followed by 12 hours in NP40 buffer (salt extraction buffer supplemented with 1% NP40 (Merck)) which allowed extraction of intracellular fractions. NP40 extracts were then analyzed on 6% Tris-Glycine gels (Thermo Fisher Scientific) under reducing conditions, followed by transfer onto nitrocellulose membranes. Membranes were then dried and exposed to a phosphoimaging plate (Fuji BAS-III or BAS-MS). After 14 days of exposure, the plates were processed using a phosphoimager (Fuji BAS 2000 or 1800), and images analyzed using ImageJ.

### Hydroxyproline Assay

Samples were incubated in 6 M hydrochloric acid (diluted in ddH_2_O, (Fluka); approximately 1 mL per 20 mg of sample) in screw top tubes (StarLab) overnight in a sand-filled heating block at 100°C, covered with aluminium foil. Tubes were then allowed to cool down and centrifuged at 12000 g for 3 min. Hydroxyproline standards were prepared (starting at 0.25 mg/mL, diluted in ddH_2_O), and serially diluted with 6 M hydrochloric acid. Each sample and standard (50 µL) were transferred into fresh Eppendorf tubes, and 450 μL of chloramine T reagent (0.127g of chloramine T in 50% N-propanol diluted with ddH_2_O; bring up to 10 mL with acetate citrate buffer – 120 g sodium acetate trihydrate, 46 g citric acid, 12 mL glacial acetic acid, 34 g sodium hydroxide, pH to 6.5 then make to 1 L with dH_2_O. All reagents from Sigma) was added to each tube and incubated at room temperature for 25 min. After, 500 μL of Ehrlich’s reagent (1.5 g of 4-dimethylamino-benzaldehyde diluted in 10 mL of N-propanol/perchloric acid at 2:1 ratio) was added to each reaction tube and incubated at 65°C for 10 minutes and then the absorbance reading at 558 nm of 100 μL of each sample was measured in a 96-well plate format.

### Mass Spectrometry

All reagents were of mass spectrometry grade. Tail tendons were homogenized with a bullet blender (with 1.6 mm steel beads; Next Advance) at maximum speed at 4°C for 5 min in 200 μL of SL-DOC buffer (1.1% SL (sodium laurate), 0.3% DOC (sodium deoxycholate), 0.5 mM dithiothreitol (DTT) in 25 mM ammonium bicarbonate) (all from Merck), supplemented with protease and phosphatase inhibitor cocktails (Roche). Samples were incubated at 4°C for 5 min, alkylated with 12 μL of 30 mM of iodoacetamide for 20 min at RT, followed by quenching with 12 μL of 30 mM DTT. Samples were centrifuged at maximum speed for 10 min. Supernatants were transferred into LoBind Protein tubes (Eppendorf) and protein concentrations measured using a Millipore Direct Detect spectrometer.

A total of 2 μg protein per sample was digested using trypsin beads (SMART digestion kit, Thermo Fisher Scientific) with rapid agitation (1400 rpm) at 37°C overnight. Peptide/bead mixtures were centrifuged at 12000 g for 5 min and supernatants transferred into new tubes. The pH was adjusted to <3 by addition of 10% formic acid (Merck) and 400 μL of EA (ethylacetate; Merck). Samples were then spun for 5 min at 16000 g, and the upper layer of solvent (EA) was discarded. EA extraction was repeated twice and samples were lyophilized in a speed-vac centrifuge (Heto).

Samples were then mixed with 100 μL of 5% acetonitrile with 0.1% formic acid (Merck), and proceeded to R3 sample desalting clean-up. Poros R3 beads (1 mg, Applied Biosystems) were added to each well of a 96-well plate with 0.2 µM PVDF membrane (Corning), and the plate was centrifuged at 200 × g for 1 min. Centrifugation was repeated to wash the beads with 50% acetonitrile (wet solution), with the flow-through discarded each time. Beads were then washed twice with wash solution (0.1% formic acid) and flow-through discarded. Samples were added into the corresponding wells and mixed with beads on a plate mixer for 5 min at 800 rpm. Samples were washed twice with wash solution, and the flow-through was discarded after each centrifugation step. A fresh sample collection plate was then placed under the PVDF membrane plate, and samples were eluted and collected using 50 μL of elution buffer (40% acetonitrile, 0.1% formic acid). This elution step was repeated to give a total of 100 μL eluted samples. Eluted samples were transferred from the plate into individual chromatography sample vials, and dried in a speed-vac centrifuge for 60 min or until complete dryness was achieved. The resulting peptides were dissolved in 15 μL of 5% acetonitrile with 0.1% formic acid.

Digested samples were analyzed using an UltiMate® 3000 Rapid Separation liquid chromatography system (RSLC, Dionex Corporation) coupled to either an Orbitrap Elite (for quality control assessment; Thermo Fisher Scientific) or Q Exactive HF (Thermo Fisher Scientific) mass spectrometer. Briefly, for the Orbitrap Elite, peptides were separated using a 75 mm × 250 μm inner diameter 1.7 μM CSH C18, analytical column (Waters) with a gradient from 92% A (0.1% formic acid, FA, Sigma, in deionized water) and 8% B (0.1% FA in acetonitrile) to 33% B, in 104 min at 300 nL/min. For the Q Exactive HF, peptides were separated in a gradient of 95% A and 5% B to 7% B at 1 min, 18% B at 58 min, 27% B in 72 min and 60% B at 74 min at 300 nL/min using a 75 mm × 250 μm inner diameter 1.7 μM CSH C18, analytical column (Waters). Peptides were selected for fragmentation automatically by data dependent analysis.

### Electron Microscopy

Tendons from 5- to 6-week old mice were prepared for transmission electron microscopy (TEM) and serial block face-scanning electron microscopy (SBF-SEM) as described previously ^70^. In brief, tendons were fixed in 1% osmium and 1.5% potassium ferrocyanide in 0.2 M sodium cacodylate buffer for 1 h, washed with distilled water, then incubated with 1% tannic acid in 0.1M cacodylate buffer for 1 h, washed with distilled water, then incubated with 1% osmium in water for 30 min. Samples were then washed with distilled water and stained with 1% uranyl acetate in water for 1 h, then dehydrated in acetone and embedded into resin.

For scanning transmission electron microscopy (STEM), mouse Achilles tendons were dissected from 6-week-old mice every 3 hours over a 24-hour period. Four mice were sacrificed at every time point. Each entire mouse Achilles was diced into ∼ 0.5 mm pieces and crushed between liquid nitrogen-cooled stainless-steel blocks. The resulting powder was then mechanically dispersed in a TRIS-EDTA buffer, pH 7.4, with added proteinase inhibitors in a hand-held Dounce homogenizer. Fibrils were adsorbed from solution on a carbon-filmed 400-mesh copper grid and washed with ultra-pure water. STEM was implemented on a Tecnai Twin (FEI, Eindhoven) electron microscope fitted with an annular dark-field detector using a camera length of 350 cm. Images were recorded at 120 kV with a sufficiently low electron dose (< 10^3^ e/nm2) to give a negligible beam-induced mass loss. Tobacco mosaic virus was used as a standard of mass per unit length.

### Mechanical Testing of Tendons

Mechanical properties of arrhythmic tendons were measured as described previously ^71^. In brief, tail tendons were dissected from 6-week-old mice and mounted onto a grade 100 sandpaper frame with a 1 cm length window using superglue. The sandpaper frame was clamped in an Instron 5943 fitted with a 10 N load cell (Instron Inc, UK). Only samples that failed in the mid portion of the tendon were included. Tendons were tested to failure with a displacement rate of 5 mm/min. The length and width of the samples tested were measured from a digital photograph using ImageJ software, which were used to derive corrected elastic modulus and calculate maximum load/cross sectional area. At least 10 tendons from each biological replicate were tested.

The dissected mouse Achilles tendons sampled at ZT3 and ZT15 were subjected to a cyclic test regime on an Instron 5943 fitted with a BioPuls specimen bath. Each sample was clamped at the bone and over the myotendinous junction using superglue and a sandpaper sandwich in the pneumatic clamps. The sample bath was filled with PBS and maintained at 22°C. The tendon was straightened with a tare load of 0.02 N and then subjected to successive strain cycles up to a 1 N load. A series of 30 cycles was used for each tendon sample and consisted of 6 sets of 5 cycles with a displacement rate for each set of 1, 2, 3, 4, 5 and 1 mm/min, respectively. The tendon was finally loaded to 1 N at 1 mm/min after the last strain cycle and the stress-relaxation load curve was recorded over 15 min. Energy loss measurements were obtained from the stress-strain hysteresis loop of the final cycle in each set. The elastic modulus was calculated from the linear part of the stress-strain curve on the load increasing part of the strain cycle. The length of the test portion of the tendon and the lateral dimensions were measured from calibrated images. The stress-relaxation curve was fitted to a sum of three exponential decay curves to yield 3 characteristic relaxation times (T1, T2, T3).

### Bioluminescence Imaging and Recordings

Tendon explants from 6-week old mice were cultured on 0.4 µm cell culture inserts (Millipore) in recording media containing 0.1 mM luciferin (Merck) in the presence of 100 nM dexamethasone (Merck) and visualized using a self-contained Olympus Luminoview LV200 microscope (Olympus) and recorded using a cooled Hamamatsu ImageEM C9100-13 EM-CCD camera. Images were taken every hour for 3 days, and combined in ImageJ to generate movies. The recording parameters were the same for control wild type (Video 6) and *Scx-Cre::Bmal1 lox/lox* (Video 7) Achilles tendons but the brightness for the *Scx-Cre::Bmal1 lox/lox* recording was increased for the images and video to enable visualization.

LumiCycle apparatus (Actimetrics) was used for real-time quantitative bioluminescence recording of tendon explants and tendon fibroblasts as described above in the absence of dexamethasone for 3 days. Then 100 nM dexamethasone was added to re-initiate the circadian rhythms. Baseline subtraction was carried out using a 24-hr moving average. Period after addition of dexamethasone was calculated using LumiCycle analysis software (Actimetrics) or using the RAP algorithm. A minimum of three peaks were used to determine period. Amplitude was calculated as peak-trough difference in bioluminescence of the second peak using base-line subtracted data.

## QUANTIFICATION AND STATISTICAL ANALYSIS

Data are presented as mean ± SEM unless otherwise indicated in figure legends. Sample number (n) indicates the number of independent biological samples in each experiment. Sample numbers and experimental repeats are indicated in figures and figure legends or methods section above. Data were analyzed using One-Way ANOVA or unpaired t-test. Data analysis was not blinded. Differences in means were considered statistically significant at p < 0.05. Significance levels are: *p<0.05; **p<0.01; ***p< 0.001; ****p< 0.0001; NS, non-significant. Analyses were performed using the Graphpad Prism 7 software.

### Morphometric Analyses of EM images

Fibril diameter, perimeter and area were measured using ImageJ software to calculate circularity and fibril volume fraction as described previously (Starborg et al., 2013). Fibril diameter distributions were plotted and fitted to 3-Gaussian distribution using Sigma Plot 12.0 software (Systat Software) as previously described (Starborg et al., 2013).

For the STEM experiments, tobacco mosaic virus (TMV) was used as standard of mass per unit length (131 kDa/nm). Mass per unit length measurements were averages of 5D-periods for each fibril. Mass per unit length measurements were made from STEM images on samples of dispersed collagen fibrils as described ^31^. The mass per unit length frequencies were weighted according to the individual fibril length in each image.

### Proteomics Data Analysis and Protein Quantification

Spectra from multiple samples were automatically aligned using Progenesis QI (Nonlinear Dynamics) with manual placement of vectors where necessary. Peak-picking sensitivity was set to 4/5 and all other parameters were left as defaults. Only peptides with charge between +1 to +4, with 2 or more isotopes were taken for further analysis. Filtered peptides were identified using Mascot (Matrix Science UK), by searching against the SwissProt and TREMBL mouse databases. The peptide database was modified to search for alkylated cysteine residues (monoisotopic mass change, 57.021 Da), oxidized methionine (15.995 Da), hydroxylation of asparagine, aspartic acid, proline or lysine (15.995 Da) and phosphorylation of serine, tyrosine, threonine, histidine or aspartate (79.966 Da). A maximum of 2 missed cleavages was allowed. Peptide detection intensities were exported from Progenesis QI as Excel (Microsoft) spread sheets for further processing. Peptide identifications were filtered via Mascot scores so that only those with a Benjamini-Hochberg false discovery rate (FDR)-adjusted p-value < 0.05 remained. FDR filtering was performed as described ^72^. Raw ion intensities from peptides belonging to proteins with fewer than 2 unique peptides (by sequence) per protein in the dataset were excluded from quantification. Remaining intensities were logged and normalized per MS run by the median logged peptide intensity. Peptides assigned to different isoforms were grouped into a single “protein group” by gene name. Only peptides observed in at least 2 samples were used in quantification. Missing values were assumed as missing due to low abundance, an assumption others have shown is justified ^73^. Imputation was performed at the peptide level following normalization using a method similar to that employed by Perseus ^74^ whereby missing values were imputed randomly from a normal distribution centered on the apparent limit of detection for this experiment. The limit of detection in this instance was determined by taking the mean of all minimum logged peptide intensities and down-shifting it by 1.6s, where s is the standard deviation of minimum logged peptide intensities. The width of this normal distribution was set to 0.3s as described ^74^. Fold-change differences in the quantity of proteins detected in different time-points were calculated by fitting a mixed-effects linear regression model for each protein with Huber weighting of residuals, as described in ^73^ using the fitglme Matlab (The MathWorks, USA) function with the formula: y_iptb= β_t+ β_p+ β_b; where y_iptb represents the log2(intensity) of peptide p belonging to protein i, at time-point t, in batch b. βs represents effect sizes for the indicated coefficients. Batch and peptide were entered as random effects whereas time-point was entered as a fixed effect. Standard error estimates were adjusted with Empirical Bayes variance correction as previously described ^75,76^. Conditions were compared with Student’s t-tests with Benjamini-Hochberg correction for false positives ^77^.

### Periodicity Analysis

Periodicity analysis was performed using the MetaCycle package ^78^ in the R computing environment ^79^ using the default parameters. A Benjamini-Hochberg FDR<0.2 was used to detect rhythmic proteins.

### Predictive Model of Collagen Homeostasis

#### Reactions and Equations

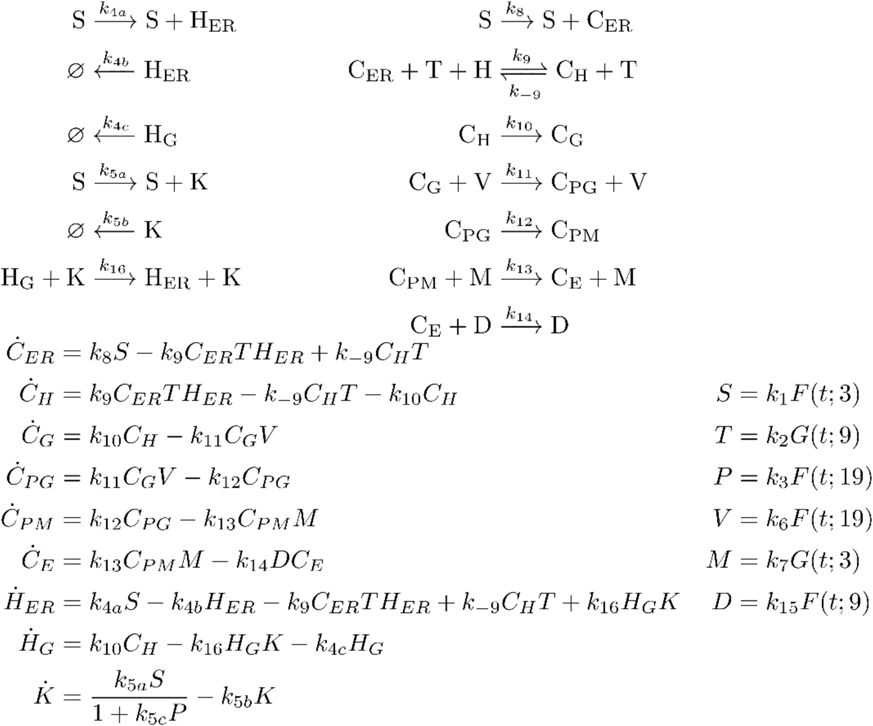

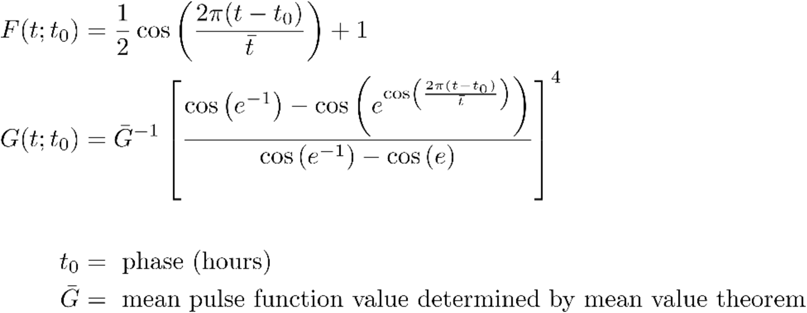

#### Variables

All variables are protein concentrations in units of *μ*M relative to typical fibroblast cell size.

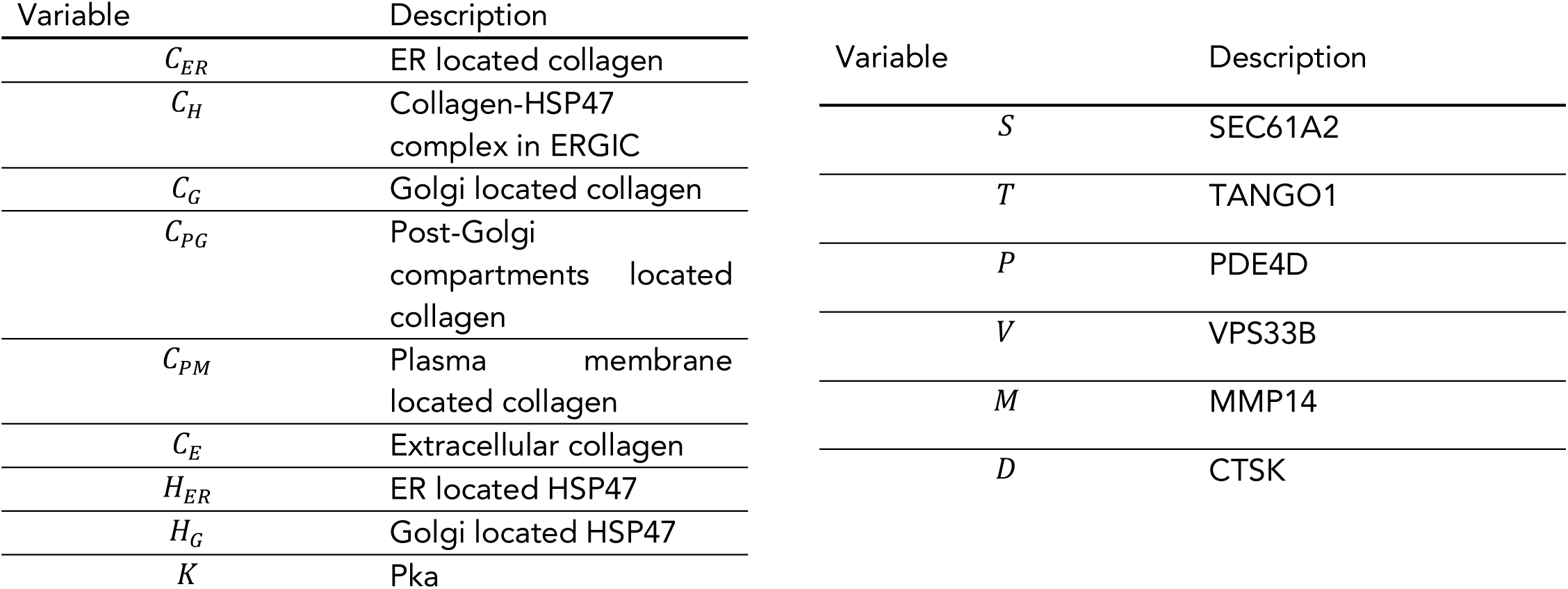

| Variable | Description |
| --- | --- |
| $C_{ER}$ | ER located collagen |
| $C_H$ | Collagen-HSP47 complex in ERGIC |
| $C_G$ | Golgi located collagen |
| $C_{PG}$ | Post-Golgi compartments located collagen |
| $C_{PM}$ | Plasma membrane located collagen |
| $C_E$ | Extracellular collagen |
| $H_{ER}$ | ER located HSP47 |
| $H_G$ | Golgi located HSP47 |
| $K$ | Pka |
| $S$ | SEC61A2 |
| $T$ | TANGO1 |
| $P$ | PDE4D |
| $V$ | VPS33B |
| $M$ | MMP14 |
| $D$ | CTSK |

#### Parameters

Parameter values for the model were derived where possible from existing results in the literature (such as ^48^). This paper lists protein copy numbers in molecules per cell, translation rates in molecules per mRNA per hour, mRNA copy numbers in molecules per cell, and protein half-life in hours, among others. These, along with a typical fibroblast cell size of 2.04 pL were combined to give translation rates in micro-molar per hour, and concentrations in micro-molar, or time scale rates in per hour. For example, *k*_1_ was calculated by dividing protein copy number by the cell size and Avogadro’s number to give concentration. Where values for a particular protein were not available, translation rates were taken as the median values from ^48^, as detailed below. The values for procollagen transport rates from one intracellular compartment to the next were guided by knowledge of the speed of vesicular transport in the eukaryotic cell.

| Parameter | Value | Description | Source |
| --- | --- | --- | --- |
| $\mu_{typ}$ | 0.0342 $\mu\text{M}$ | Median typical protein concentration | Median value calculated from <sup>48</sup> |
| $k_1$ | 0.00822 $\mu\text{M}$ | SEC61A2 mean protein concentration | Calculated directly from <sup>48</sup> |
| $k_2$ | 0.0105 $\mu\text{M}$ | TANGO1 mean protein concentration | Calculated directly from <sup>48</sup> |
| $k_3$ | 0.0342 $\mu\text{M}$ | PDE4D mean protein concentration | Median value calculated from <sup>48</sup> |
| $k_6$ | 0.00335 $\mu\text{M}$ | VPS33B mean protein concentration | Calculated directly from <sup>48</sup> |
| $k_7$ | 0.00457 $\mu\text{M}$ | MMP14 mean protein concentration | Calculated directly from <sup>48</sup> |
| $k_{15}$ | 0.0342 $\mu\text{M}$ | CTSK mean protein concentration | Median value calculated from <sup>48</sup> |
| $\bar{t}$ | 24 h | Circadian period | Length of one circadian cycle |
| $k_{4a}$ | 4.84 $\text{h}^{-1}$ | HSP47 protein synthesis rate | Adjusted for physicality from values calculated directly from <sup>48</sup> |
| $k_{4b}$ | 0.00413 $\text{h}^{-1}$ | ER HSP47 protein degradation rate | Calculated directly from <sup>48</sup> |
| $k_{4c}$ | 0.00413 $\text{h}^{-1}$ | Golgi HSP47 protein degradation rate | Calculated directly from <sup>48</sup> |
| $k_{5a}$ | 1.89 $\text{h}^{-1}$ | PKA protein synthesis rate | Calculated directly from <sup>48</sup> |
| $k_{5b}$ | 0.00690 $\text{h}^{-1}$ | PKA protein degradation rate | Calculated directly from <sup>48</sup> |
| $k_8$ | 0.0693 $\text{h}^{-1}$ | Col-I synthesis rate | Adjusted for physicality from values calculated directly from <sup>48</sup> |
| $k_{10}$ | 3600 $\text{h}^{-1}$ | ERGIC to Golgi Col-I transition rate | Estimated time scale (~1 second) of ERGIC vesicular transport calculated from typical cellular vesicle speeds and inter-compartmental distances <sup>80</sup> |
| $k_{12}$ | 0.0417 $\text{h}^{-1}$ | Post-Golgi compartments to plasma membrane Col-I transition rate | Estimated from dimensionally consistent powers of $\bar{t}$ |
| $k_{5c}$ | 29.2 $\mu\text{M}^{-1}$ | Rate constant for repression of PKA by PDE4D | Estimated from dimensionally consistent products of $\mu_{typ}$ and $\bar{t}$ |
| $k_{-9}$ | 1.22 $\mu\text{M}^{-1}\text{h}^{-1}$ | Col-I-HSP47 dissociation rate | Estimated from dimensionally consistent products of $\mu_{typ}$ and $\bar{t}$ |
| $k_{11}$ | 1.22 $\mu\text{M}^{-1}\text{h}^{-1}$ | Golgi to Post-Golgi | Estimated from dimensionally consistent |

| | | compartments Col-I transition rate | products of $\mu_{typ}$ and $\bar{t}$ |
| --- | --- | --- | --- |
| $k_{13}$ | $12.2 \mu\text{M}^{-1}\text{h}^{-1}$ | Plasma membrane to extracellular Col-I transition rate | Estimated from dimensionally consistent products of $\mu_{typ}$ and $\bar{t}$ |
| $k_{14}$ | $1.22 \mu\text{M}^{-1}\text{h}^{-1}$ | CTSK dependant extracellular Col-I degradation rate | Estimated from dimensionally consistent products of $\mu_{typ}$ and $\bar{t}$ |
| $k_{16}$ | $1.22 \mu\text{M}^{-1}\text{h}^{-1}$ | Golgi to ER HSP47 transition rate | Estimated from dimensionally consistent products of $\mu_{typ}$ and $\bar{t}$ |
| $k_9$ | $0.356 \mu\text{M}^{-2}\text{h}^{-1}$ | Col-I-HSP47 complex association rate | Estimated from dimensionally consistent products of $\mu_{typ}$ and $\bar{t}$ |

#### Collagen Fiber Quantification

Collagen fibers were independently and manually scored by two independent researchers using a Zeiss fluorescence microscope. An average of at least 12 fields of views were quantified for each time point. Number of fibers were counted based on collagen-I fluorescence staining; where collagen fibers appear branched it was counted as separate fibers. For each field of view, the number of fibers were corrected to the number of cells as defined by DAPI staining. For each time point, the number of fibers per cell was corrected to the time point with the maximum value, expressing all other quantifications as a percentage of the maximum. Representative images from selected time points were presented.

## DATA AND SOFTWARE AVAILABILITY

The mouse tail tendon time-series microarray data ^29^ have been deposited to Array Express repository with the dataset identifier (ArrayExpress accession number E-MTAB-7743). All raw mass spectrometric data files from this study are available for download in Supplementary Information.

## Acknowledgements

The research was funded by Wellcome (110126/Z/15/Z and 203128/Z/16/Z), MRC (MR/P010709/1) and Arthritis Research UK (20875). VM was supported by a studentship from the Sir Richard Stapley Educational Trust. JS was funded by a Biotechnology and Biological Sciences Research Council (BBSRC) David Phillips Fellowship (BB/L024551/1). BC is supported by a Wellcome 4-year PhD studentship (210062/Z/17/Z). The authors would like to thank Ronen Schweitzer (Shriners Hospital for Children, Portland, OR, USA) for *Scx-Cre* mice, Alison Hallworth and Raymond Hodgkiss and the University of Manchester Biological Support Facility for assistance in animal welfare and husbandry, and Michal Dudek and Nan Yang (University of Manchester) for assistance with bioluminescence imaging. The proteomics was performed at the Biological Mass Spectrometry Facility in the Faculty of Biology, Medicine and Health (University of Manchester) with the assistance of Stacey Warwood and Ronan O’Cualain, and electron microscopy was performed in the Electron Microscopy Facility, Faculty of Biology, Medicine and Health (University of Manchester).

## Author contributions

C-YCY and RG designed and performed experiments, interpreted data and drafted the manuscript. AP, JC, DFH, YL and BC designed and performed experiments and interpreted data. VM, JS and AA contributed materials and analysis tools. HR-H, OJ and TS supervised the mathematical aspects of the research. Q-JM and KEK conceived the project, supervised the experiments and wrote the manuscript.

## Figure legends

**Supplementary Figure 1**

**(A)** Collagen fibril diameter distributions measured from transverse TEM images of tail tendons sampled through a circadian period. Mice were housed in 12-hour dark/12-hour light cycles. ZT, zeitgeber time (hours into light). Greater than 1000 fibrils measured in each panel. Black lines show 3-Gaussian fit curves. **(B)** Typical scanning transmission electron microscopy image of fibrils from mechanically disrupted tendon. Bar, 500 nm. **(C)** Representative mass per unit length distribution measured from scanning transmission electron microscopy images of mechanically disrupted whole tendons. Time point shown is ZT12. Percentage of D1 fibrils at all time points over 24 h is shown in Figure 1E. **(D)** Elastic modulus of Achilles tendon is not time-of-day dependent (n=4). **(E)** Representative hysteresis curves from cyclic loading show the energy loss is greater in tendons taken at ZT15 compared with ZT3. **(F)** Time constants derived from stress-relaxation from an initial 1 N load of Achilles tendon sampled at ZT3 compared to ZT15 (T_1_, T_2_ and T_3_) (n=4; *p<0.05). **(G)** Comparison of energy loss values of Achilles tendons at ZT3 versus ZT15 for a 5-fold range of strain rates showing a consistent ∼ 50% greater energy loss at ZT15. (n=4, **p<0.01, ***p<0.001).

**Supplementary Figure 2**

Expression of collagen genes: **(A)** *Col1a1, Col1a2*, **(B)** *Col2a1, Col3a1*, **(C)** *Col5a1, Col5a2, Col5a3*, **(D)** *Col11a1, Col11a2*, in wild type mouse tail tendon tissue detected by different probe sets in a time series microarray study were not rhythmic (n=2; bars show SEM). Grey shadow indicates subjective night phase.

**Supplementary Figure 3**

**(A)** Images of electron microscopy of transverse sections across mouse tendons fibroblasts *in vivo* showing the endoplasmic reticulum (ER) at different time points. Bars 500 nm. **(B)** Quantification of ribosome docking onto ER show differences at ZT2 and ZT15.5 in tendon fibroblasts *in vivo* (***p<0.001). **(C)** Relative expression of *Sec61a1* and *Sec61a2* mRNA was reduced in MEFs after CRISPR-Cas9-mediated deletion of both *Sec61a1* and *Sec61a2* compared to control wild type MEFs (n=3; *p<0.05, ***p<0.001). **(D)** Immunofluorescence analysis of collagen-I entry into the ER (detected via the marker calnexin) showed little co-localization of collagen-I and calnexin in *Sec61a1* and *Sec61a2* KO MEFs compared to wild type MEFs. **(E)** Relative expression of *Mia3* mRNA was reduced in MEFs treated with siRNA targeting *Mia3* compared to cells treated with scrambled control siRNA (n=3; ***p<0.001). **(F)** Relative expression of *Pde4d* mRNA was reduced in MEFs treated with siRNA targeting *Pde4d* compared to cells treated with scrambled control siRNA (n=3; ***p<0.001). **(G)** Western blot analysis of MEFs treated with siRNAs targeting *Pde4d* showed depletion of PDE4D protein (*Pde4d* KD). Levels of GAPDH protein served as a loading control. **(H)** Relative expression of *Vps33b* mRNA was significantly reduced in MEFs treated with CRISPR/Cas9n targeting *Vps33b* compared to control. Controls were wild type MEFs that underwent electroporation with Cas9 protein but without gRNA (n=2; ***p<0.001). **(I)** Relative expression of *Vps33b* mRNA in iTTFs after CRISPR/Cas9-mediated deletion of *Vps33b* (n=3; **p<0.01). **(J)** Western blot analysis of VPS33b protein from *Vps33b* KO immortalized tail tendon fibroblasts (iTTFs). Vinculin protein served as a loading control.

**Supplementary Figure 4**

MEFs were synchronized and collagen-I deposition was assessed by indirect immunofluorescence every 4 hours during 60 hours. Representative images from each time point are shown. Culture of synchronized cells was cultured in the presence of 1 mM DMOG as a control for no collagen secretion. Bars 10 µm.

**Supplementary Figure 5**

MEFs were synchronized and collagen-I co-localization with GM130 was assessed by indirect immunofluorescence every 4 hours during 24 hours (up to 15 hours post-synchronization shown). Arrows point to the Golgi. Bars 10 µm.

**Supplementary Figure 6**

(**A**) TEM of mouse tail tendons prepared in PBS. (**B**) TEM of mouse tail tendons after disruption in SLDOC and collection by centrifugation. (**C**) Diameter distribution of collagen fibrils in unextracted tendon (untreated) and after extraction by SLDOC (pellet). (**D**) Hydroxyproline content of dissected tail tendons from mice incubated with SL-DOC. Data are expressed in percentage of total collagen extracted from tendons demonstrating that SL-DOC extracts 10-15% of total collagen in the tendon (n=2; standard deviation shown).

**Supplementary Figure 7**

**(A)** The 141 rhythmic proteins were separated into CT1-12 (day) and CT13-24 (night). The numbers of proteins in each group is shown. These two lists were analyzed using ENRICHR (27141961). **(B)** Gene ontology terms based on molecular function for each group are shown. **(C, D)** Achilles and tail tendon tissues were harvested at the day phase (ZT3, day) or night phase (ZT15, night) and labeled with ^14^C-proline for 1 h in the presence of exogenous L-ascorbate. Higher incorporation of ^14^C was observed in the night phase in both Achilles (n=5; *p=0.0423) and tail tendons (n=3; *p=0.0395). **(E)** Phosphoimager analysis of 1 h ^14^C-proline labeling of mouse Achilles tendon. Proteins were extracted into neutral buffer containing NP40 buffer extraction. Higher incorporation of ^14^C is observed in the night phase. **(F)** Western blot analysis of the samples shown in **(E)**. Collagen-I was observed with an anti-collagen-I antibody.

**Supplementary Figure 8**

**(A)** Luminometry recordings of endogenous circadian rhythms and re-initiation of the rhythms with dexamethasone (arrow) evident in dissected Achilles and tail tendons of wild type mice but not of *Scx-Cre::Bmal1 lox/lox* mice bred on a *PER2::Luciferase* background. A residual rhythm observed in the *Scx-Cre*::*Bmal1 lox/lox* tendons was likely due to cells that were not derived from Scleraxis-lineage cells. Relative amplitude and period of wild type and *Scx-Cre::Bmal1 lox/lox* tail and Achilles tendon circadian rhythms at 6-weeks of age (n=5 tail tendons, n=4 Achilles tendons; *p<0.05, **p<0.001). **(B)** Representative transverse transmission electron microscopy images of tail tendons from 6 week-old *Scx-Cre*::*Bmal1 lox/lox* mice showing irregular fibril cross sections. Bars 200 nm. **(C)** Fibril diameter distributions measured from transverse TEM images of *Scx-Cre::Bmal1 lox/lox* tail tendons showing abnormal distribution of fibril diameters and increase in large (>400-nm fibrils) in mutant tendons (>1000 fibrils measured). Red lines show 3-Gaussian fit curves. **(D)** Circularity of D1, D2 and D3 collagen fibrils of tail tendons sampled from *Scx-Cre::Bmal1 lox/lox* mice and their wild type litter mate controls (664 wild type fibrils and 506 *Scx-Cre::Bmal1 lox/lox* fibrils measured; ***p<0.0001 in D1, D2 and D3 populations in mutants compared to same populations in littermate controls). The circularity of fibrils from global *ClockΔ19* tendons (Figure 7K) being more irregular than fibrils from *Scx-Cre::Bmal1 lox/lox* is likely due to due to a residual circadian rhythm from cells that were not derived from Scleraxis-lineage cells. **(E)** *ClockΔ19* tendons are mechanically weaker (n=6; *p<0.05, **p<0.01) and thicker (n=6; **p<0.01) than age-matched (6-weeks old) wild type mice. Fibril volume fraction is unchanged in *ClockΔ19* tendons compared to wild type tendons. **(F)** *Scx-Cre:Bmal1 lox/lox* tendons are mechanically weaker (n=6; **p<0.01, ***p <0.001) and thicker (n=6; **p<0.01) than age-matched (6-weeks old) wild type mice.

## Supplementary Data

MetaCycle output (columns A-X) for timed series LC-tandem mass spectrometry analysis of proteins extracted from 6-week-old mouse male tail tendon (columns Y-AJ).

### Video 1

Step-through movie of serial block face-scanning electron microscopy of mouse Achilles tendon at 09:00 hrs. Bar, 7 µm. Z-depth, 486 µm, images recorded at 9.1 µm intervals to reduce file size.

### Video 2

Step through movie displaying individual colourised fibrils in small region of mouse tail tendon, showing merging of D1 fibrils onto larger D2 and D3 fibrils. Data gathered from serial block face-scanning EM stack of images. Area shown is 4 × 4 µm, each frame is a longitudinal step of 100 nm through the tendon.

### Video 3

Step-through movie of serial block face-scanning EM of mouse Achilles tendon at 09:00 hrs. Bar, 500 nm. D3 fibril highlighted with a blue line. Filled circles indicate D1 fibrils.

### Video 4

Step-through movie of serial block face-scanning electron microscopy of mouse Achilles tendon showing diameter variability along the length of fibrils and tortuosity of D1 fibrils. Bar, 700 nm. Z-depth, 12.5 µm.

### Video 5

Step-through animation showing the relative proportion of PC-I in the ER, ER exit sites, Golgi, post-Golgi, plasma membrane, and extracellular matrix, starting at CT0.

### Video 6

Real-time bioluminescence microscopy of dissected Achilles tendon from wild type *PER2::Luciferase* mice during 5 days.

### Video 7

Real-time bioluminescence microscopy of dissected Achilles tendon from *ScxCre::Bmal1* lox/lox *PER2::Luciferase* mice during 5 days. The brightness for the *Scx-Cre::Bmal1* lox/lox tendon recording was increased to enable visualization.

## KEY RESOURCES TABLE

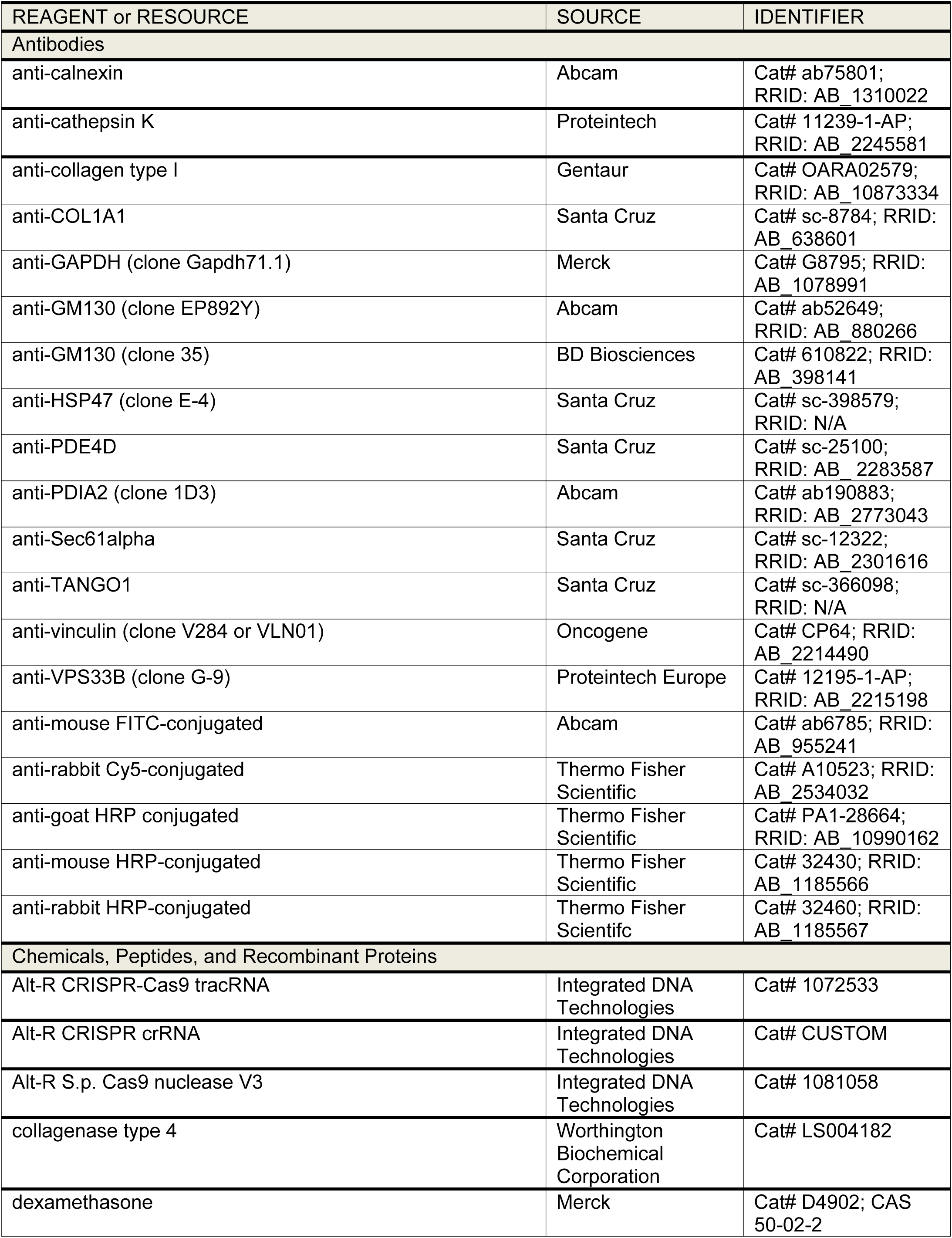
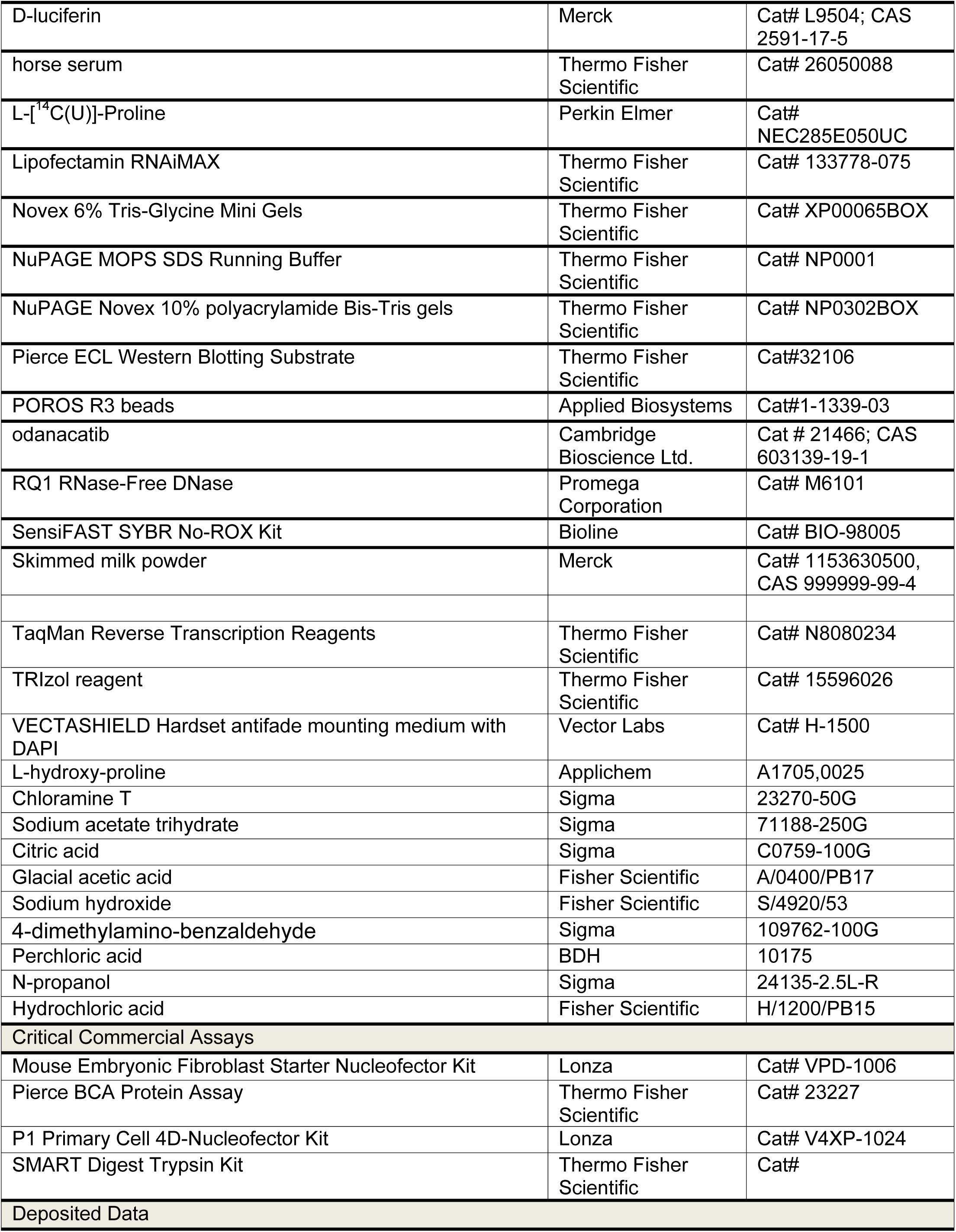
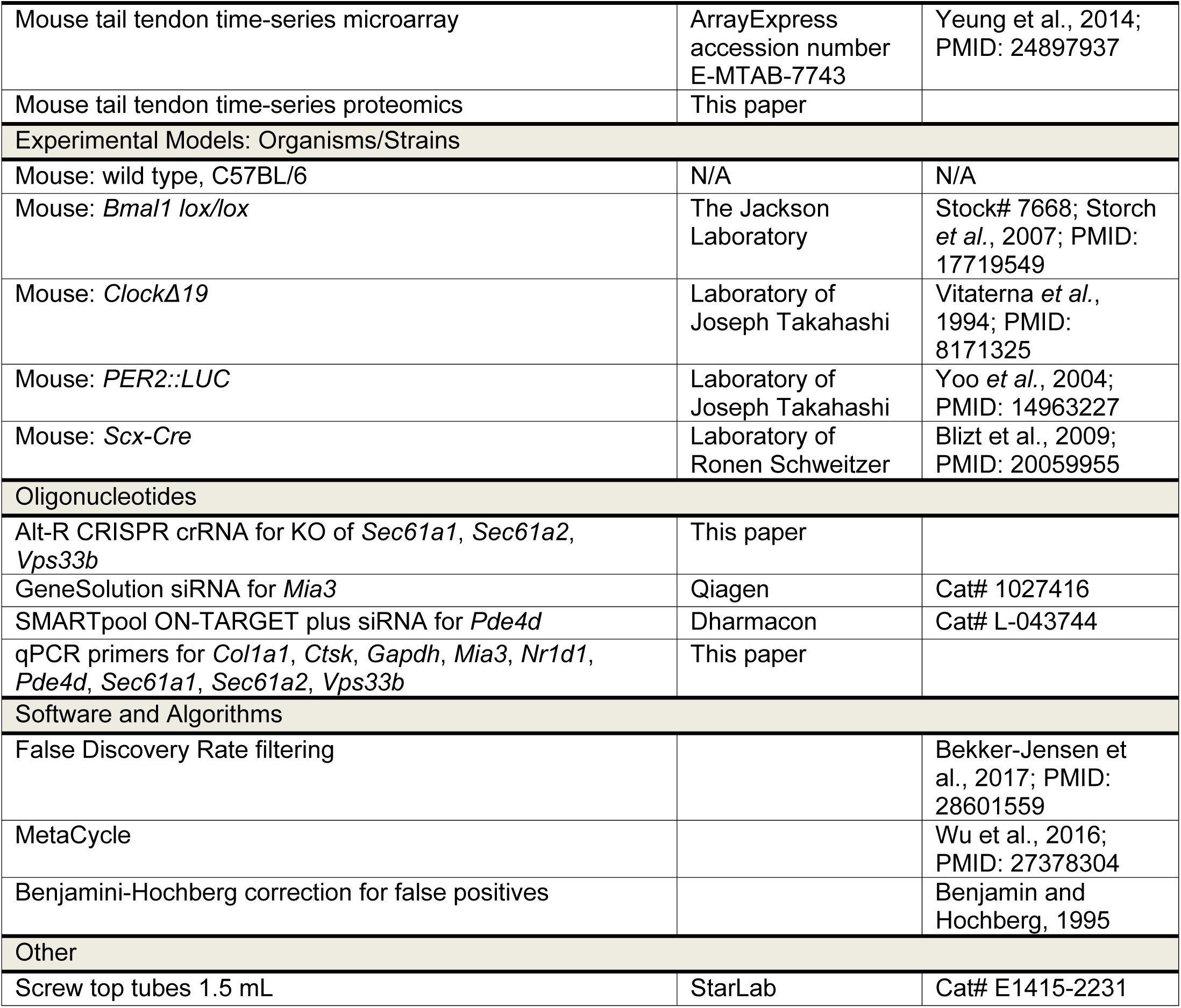

