## Supplementary figures for "Circadian control of the secretory pathway is a central mechanism in tissue homeostasis"

Chang, Garva, Pickard, Yeung et al. – Supplementary Figure 1

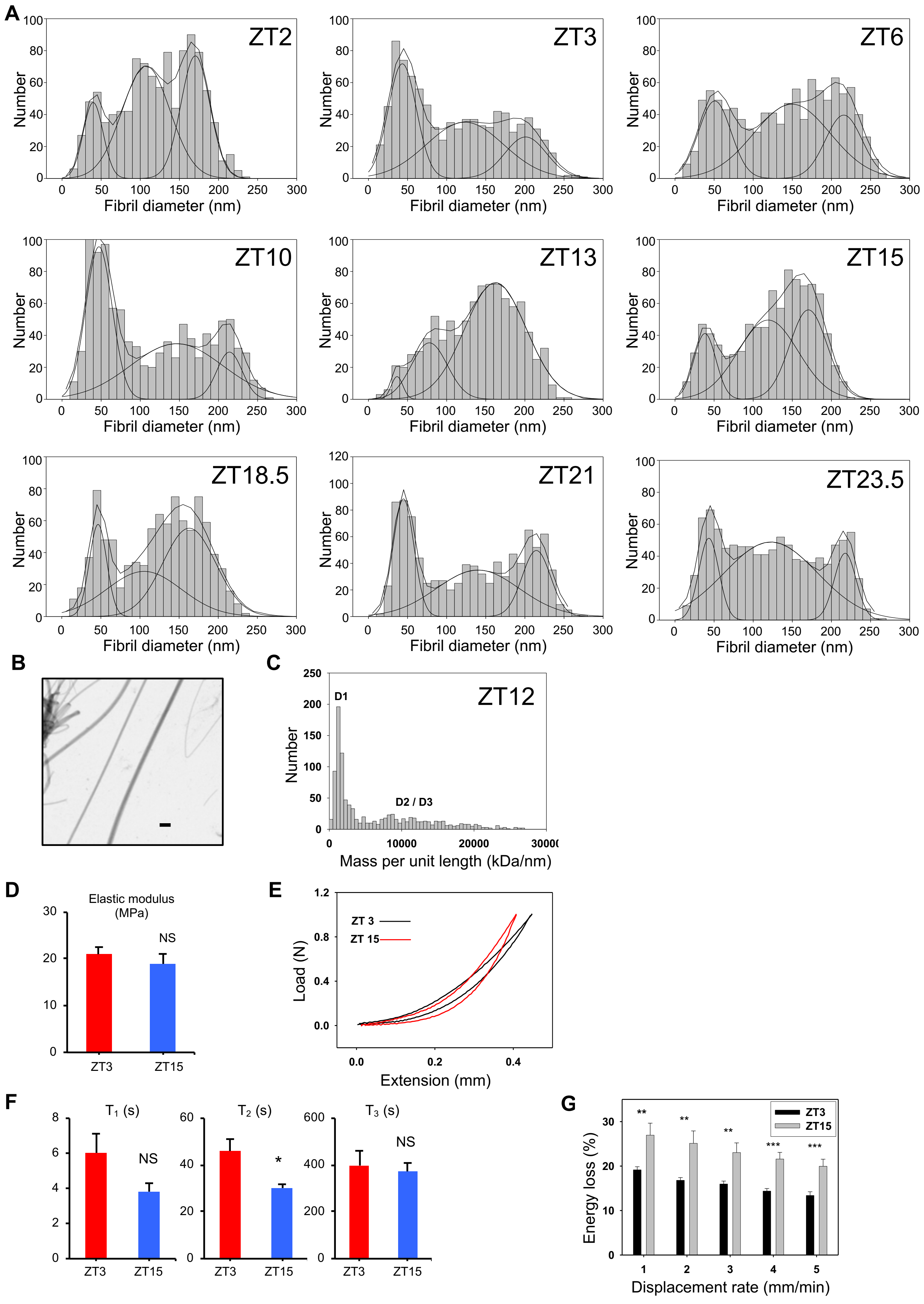

Chang, Garva, Pickard, Yeung et al. – Supplementary Figure 2

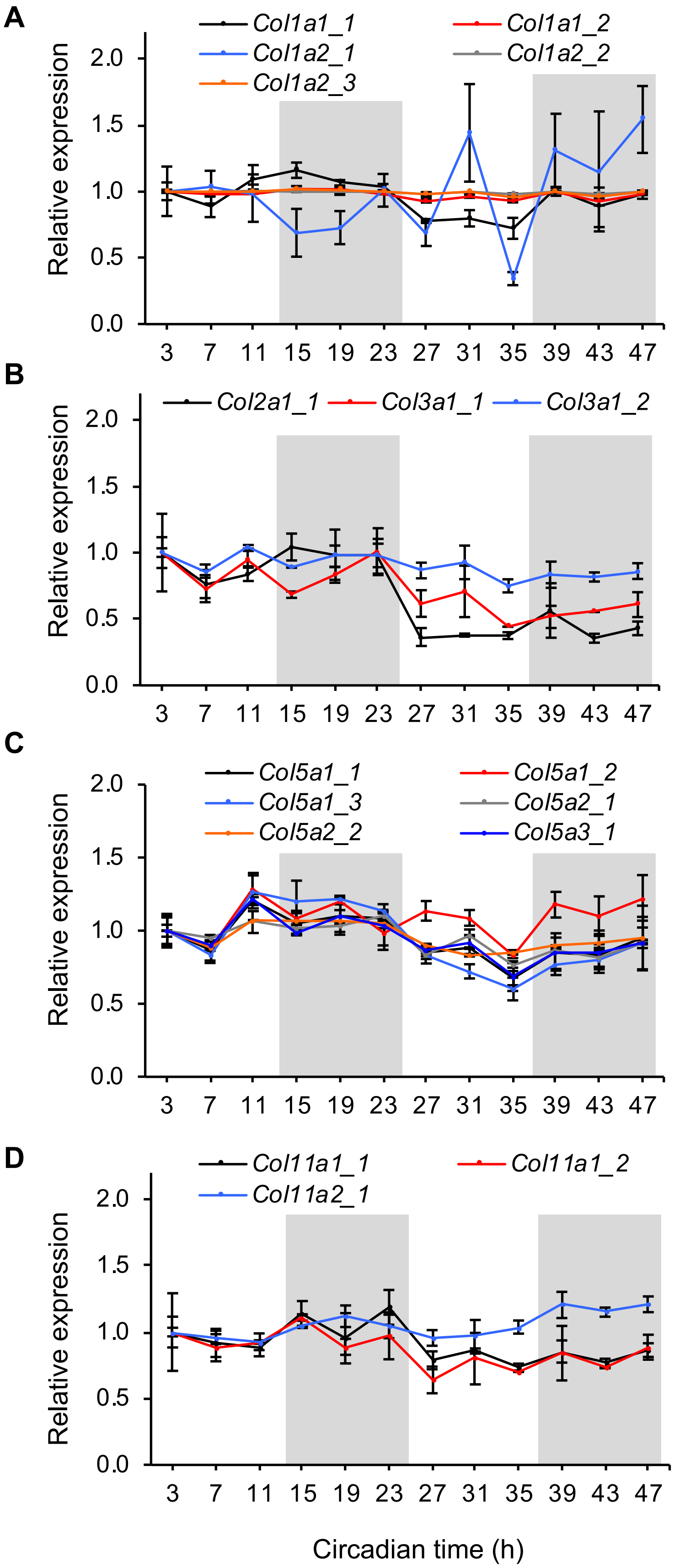

Chang, Garva, Pickard, Yeung et al. – Supplementary Figure 3

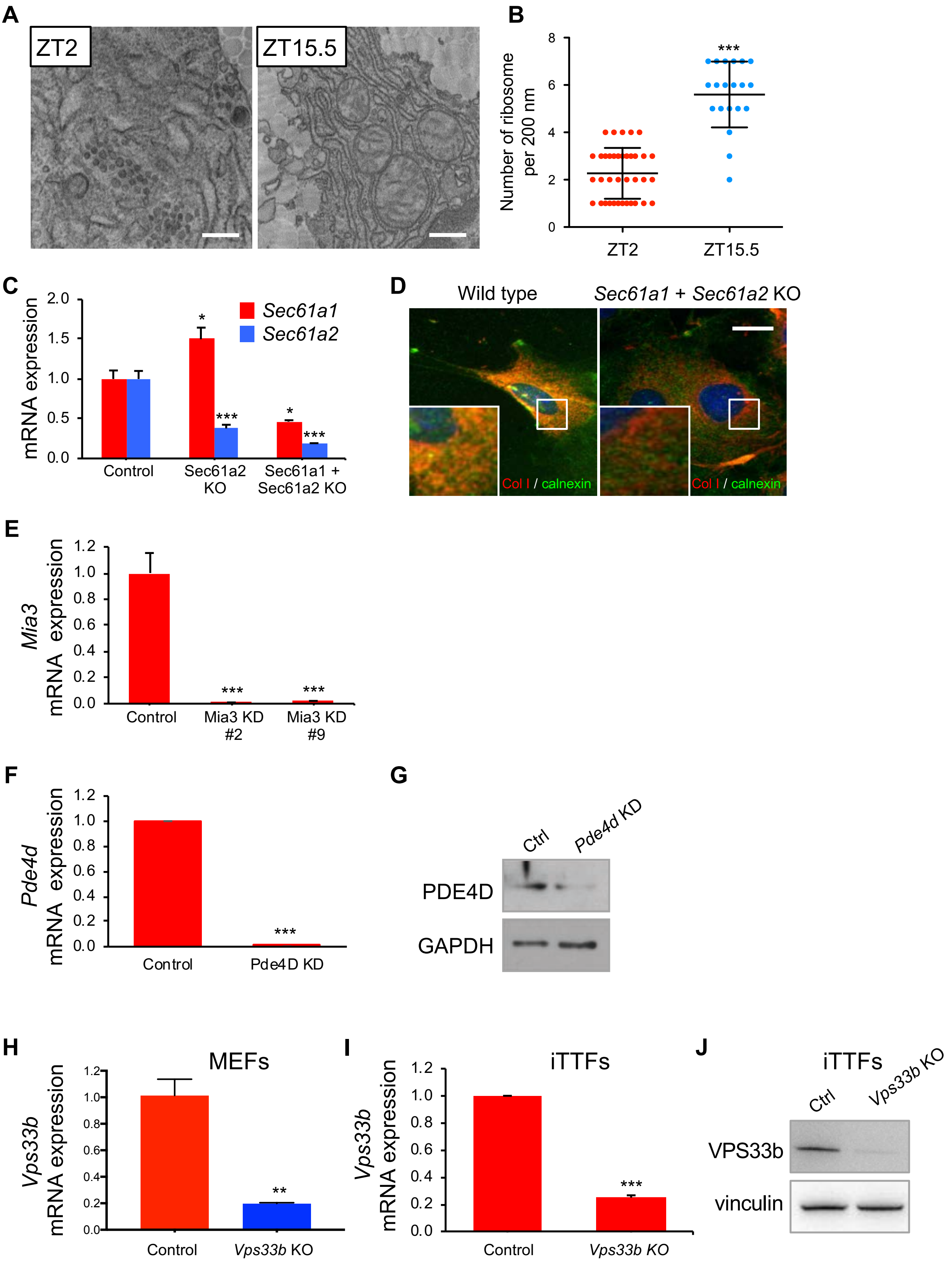

### Chang, Garva, Pickard, Yeung et al. – Supplementary Figure 4

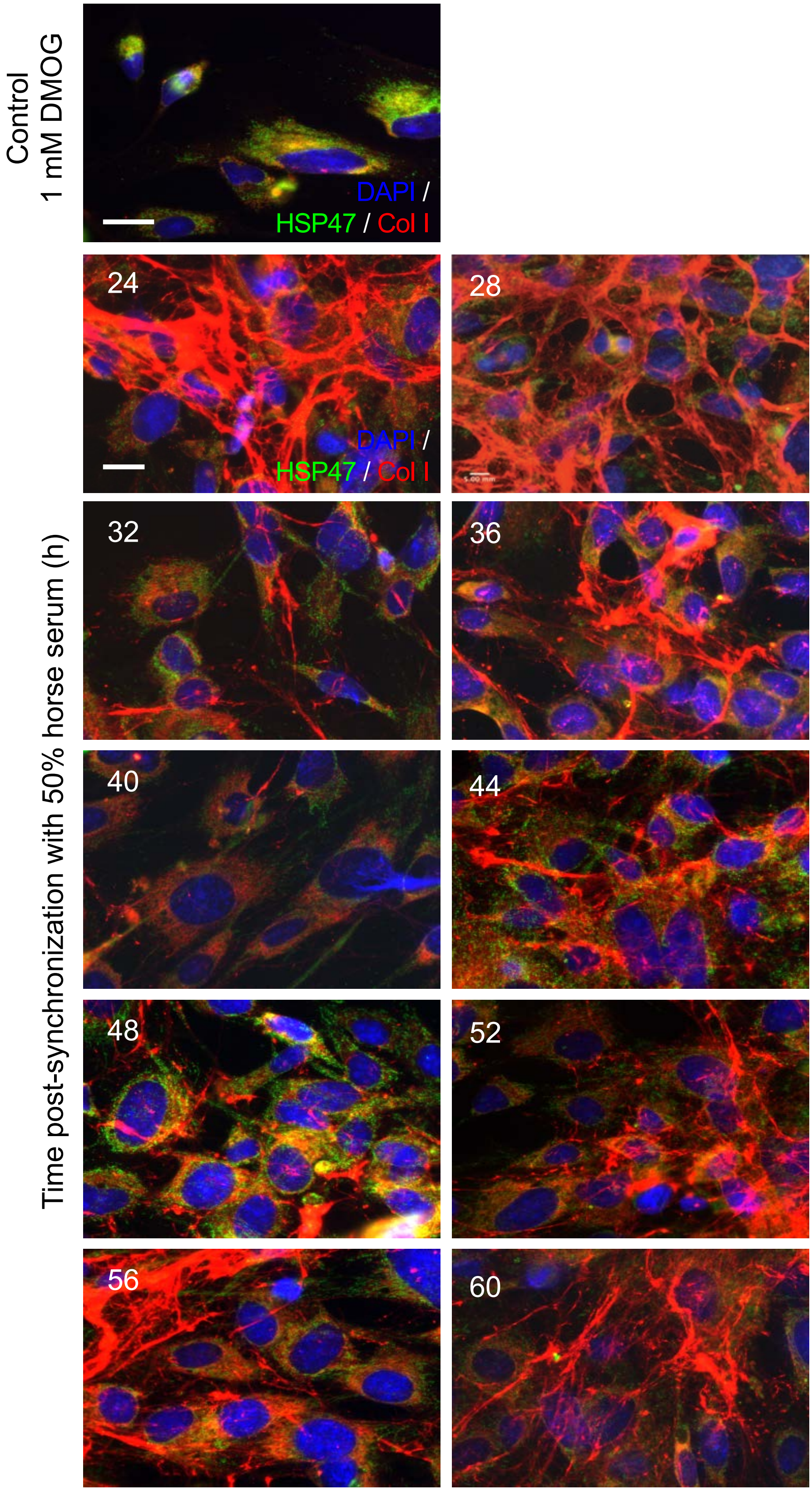

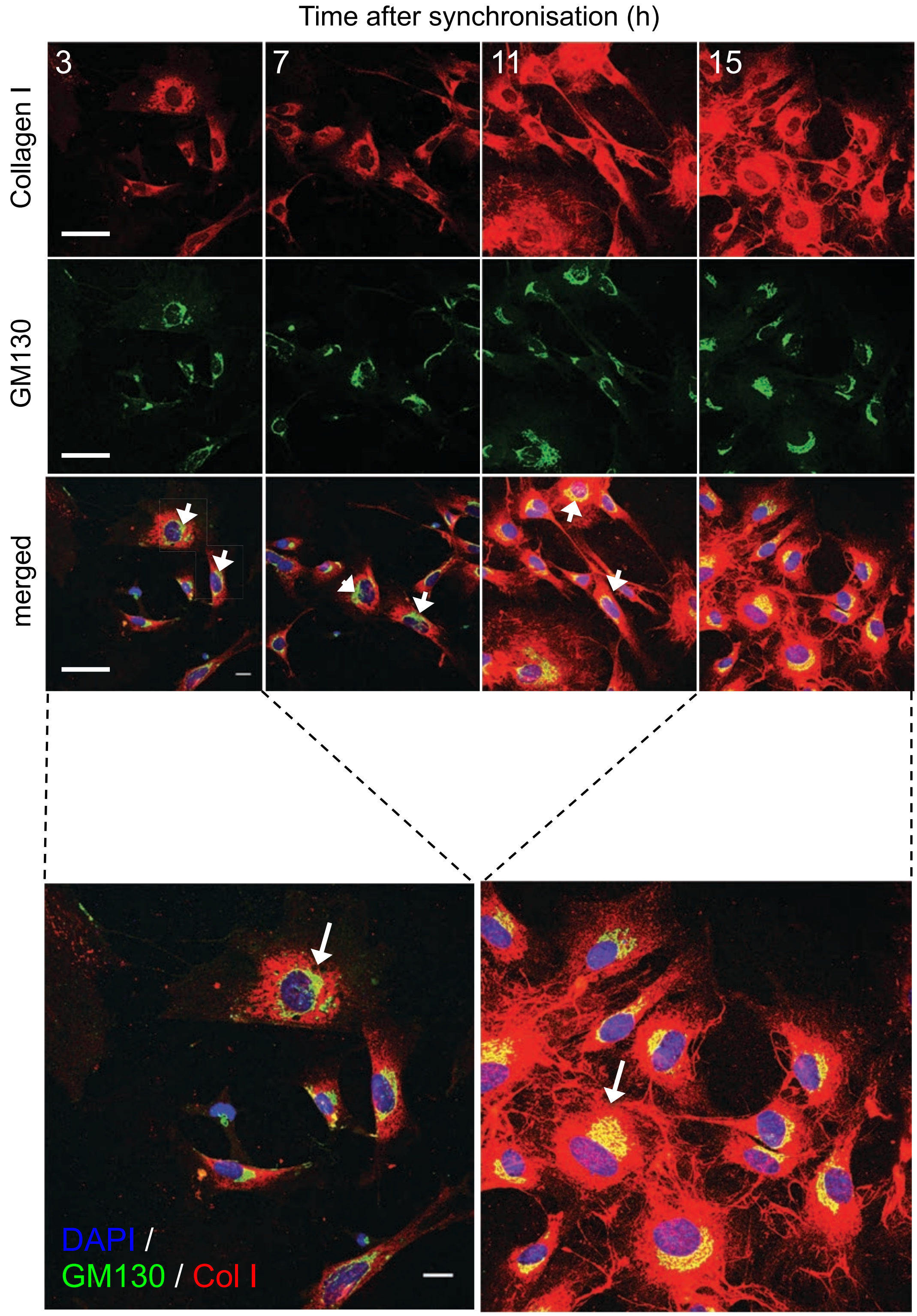
Time after synchronisation (h)
3
7
11
15
Collagen I
GM130
merged
DAPI /
GM130 / Col I

**A**

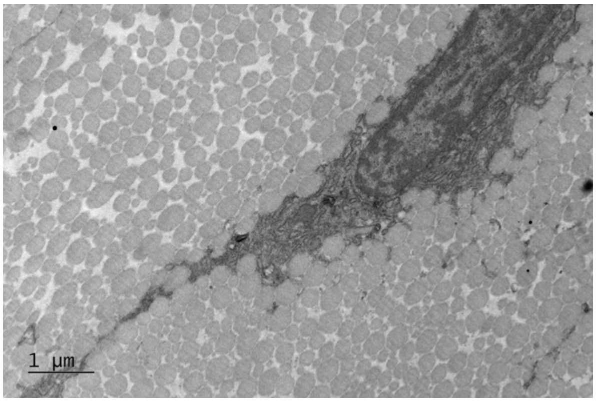
1 µm

**B**

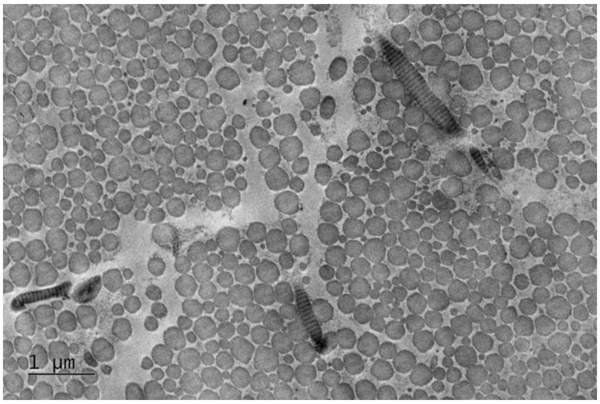
1 µm

**C**

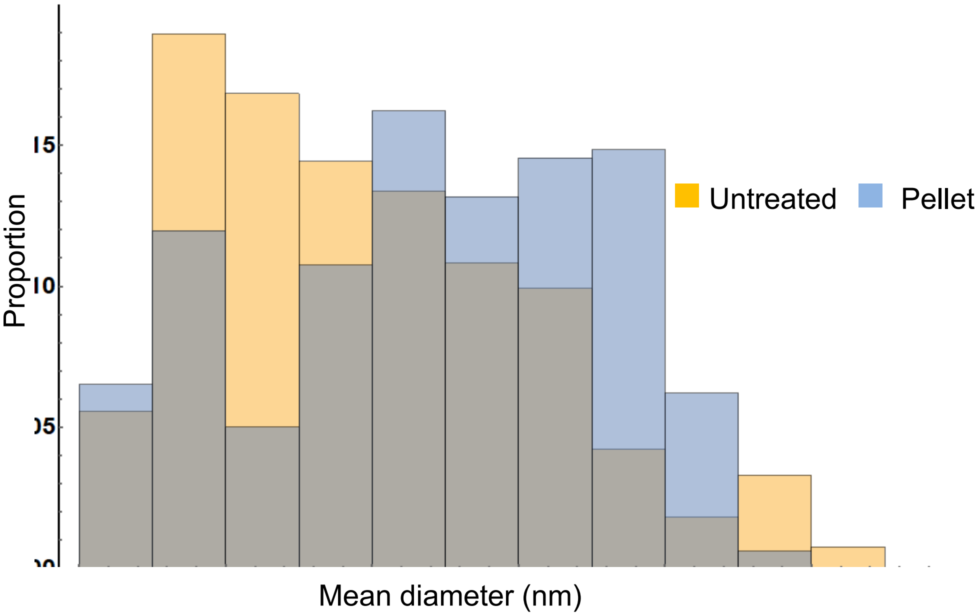
Proportion
Untreated
Pellet
Mean diameter (nm)

**D**

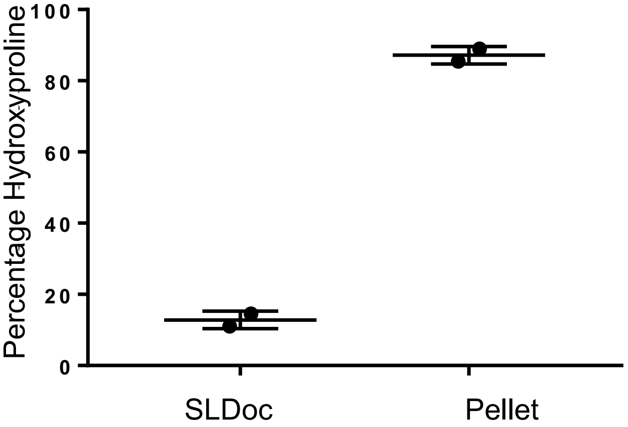
Percentage Hydroxyproline
100
80
60
40
20
0
SLDoc
Pellet

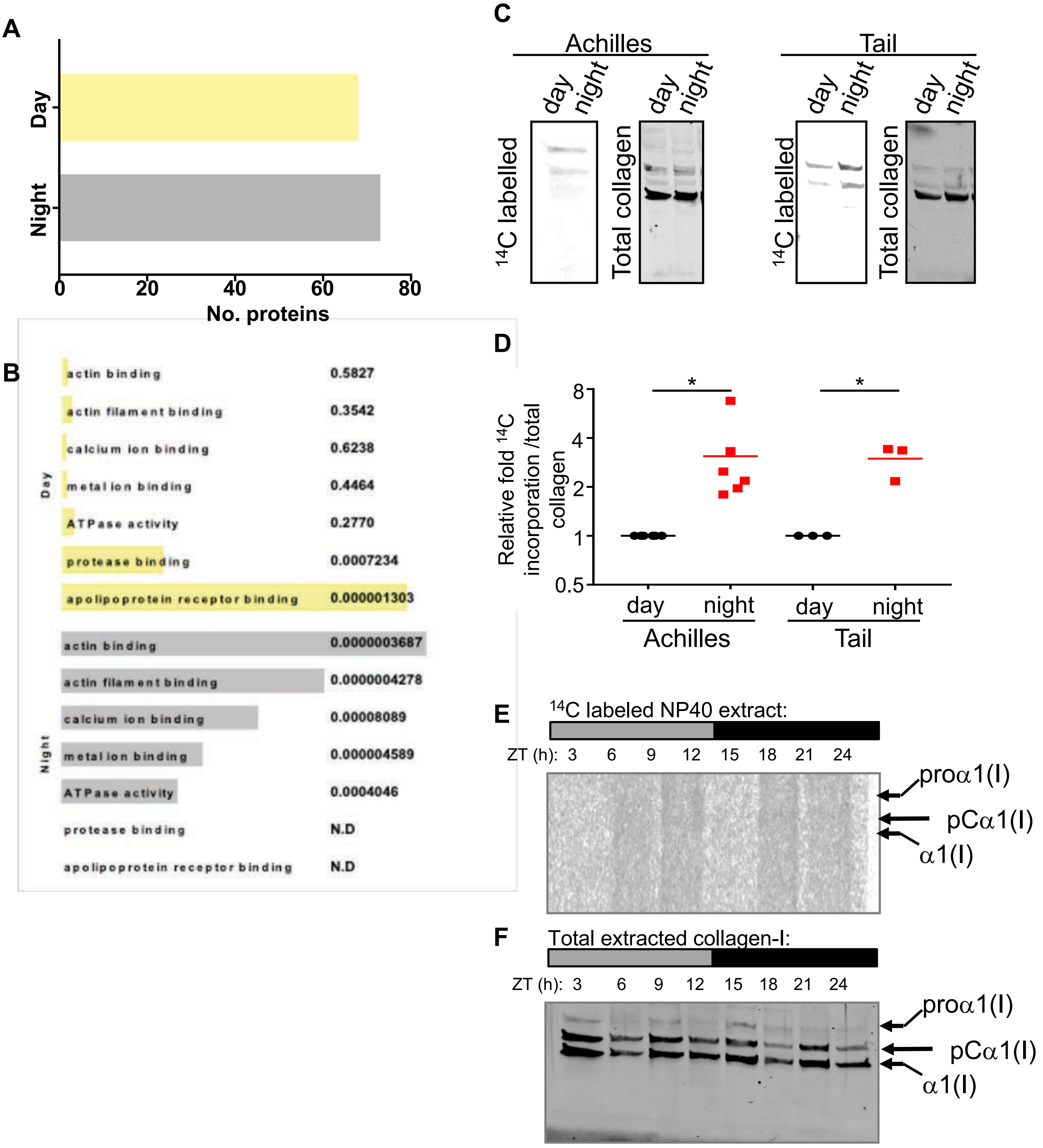

A
Day
Night
0
20
40
60
80
No. proteins
B
Day
actin binding 0.5827
actin filament binding 0.3542
calcium ion binding 0.6238
metal ion binding 0.4464
ATPase activity 0.2770
protease binding 0.0007234
apolipoprotein receptor binding 0.000001303
Night
actin binding 0.0000003687
actin filament binding 0.0000004278
calcium ion binding 0.00008089
metal ion binding 0.000004589
ATPase activity 0.0004046
protease binding N.D
apolipoprotein receptor binding N.D
C
Achilles
Tail
day night
day night
14C labelled
Total collagen
D
Relative fold 14C incorporation /total collagen
8
4
2
1
0.5
*
day night
Achilles
day night
Tail
E
14C labeled NP40 extract:
ZT (h): 3 6 9 12 15 18 21 24
proα1(I)
pCα1(I)
α1(I)
F
Total extracted collagen-I:
ZT (h): 3 6 9 12 15 18 21 24
proα1(I)
pCα1(I)
α1(I)

Chang, Garva, Pickard, Yeung et al. – Supplementary Figure 8

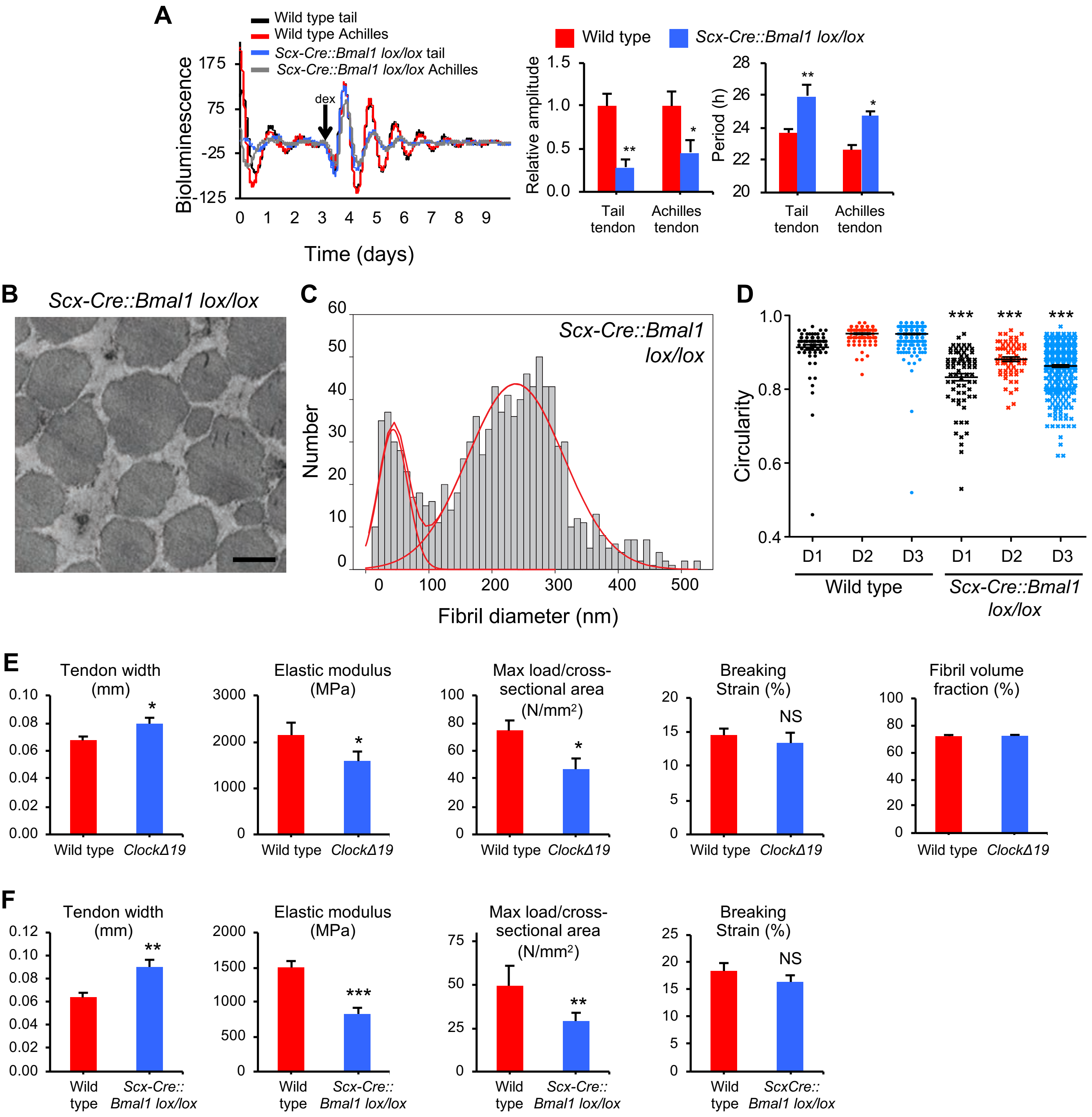
